# Dissecting the heterogeneous cortical anatomy of autism spectrum disorder using normative models

**DOI:** 10.1101/477596

**Authors:** Mariam Zabihi, Marianne Oldehinkel, Thomas Wolfers, Vincent Frouin, David Goyard, Eva Loth, Tony Charman, Julian Tillmann, Tobias Banaschewski, Guillaume Dumas, Rosemary Holt, Simon Baron-Cohen, Sarah Durston, Sven Bölte, Declan Murphy, Christine Ecker, Jan K. Buitelaar, Christian F. Beckmann, Andre F. Marquand

## Abstract

**Background:** The neuroanatomical basis of autism spectrum disorder (ASD) has remained elusive, mostly due to high biological and clinical heterogeneity among diagnosed individuals. Despite considerable effort towards understanding ASD using neuroimaging biomarkers, heterogeneity remains a barrier, partly because studies mostly employ case-control approaches, which assume that the clinical group is homogeneous.

**Methods:** Here, we used an innovative normative modelling approach to parse biological heterogeneity in ASD. We aimed to dissect the neuroanatomy of ASD by mapping the deviations from a typical pattern of neuroanatomical development at the level of the individual and to show the necessity to look beyond the case-control paradigm to understand the neurobiology of ASD. We first estimated a vertex-wise normative model of cortical thickness development using Gaussian process regression, then mapped the deviation of each participant from the typical pattern. For this we employed a heterogeneous cross-sectional sample of 206 typically developing (TD) individuals (127 male), and 321 individuals (232 male) with ASD (aged 6-31).

**Results:** We found few case-control differences but the ASD cohort showed highly individualized patterns of deviations in cortical thickness that were widespread across the brain. These deviations correlated with severity of repetitive behaviors and social communicative symptoms, although only repetitive behaviors survived corrections for multiple testing.

**Conclusions:** Our results: (i) reinforce the notion that individuals with ASD show distinct, highly individualized trajectories of brain development and (ii) show that by focusing on common effects (i.e. the ‘average ASD participant’), the case-control approach disguises considerable inter-individual variation crucial for precision medicine.

## Introduction

Autism spectrum disorder (ASD) is a lifelong neurodevelopmental disorder diagnosed exclusively on the basis of symptomatology, period of onset, and impairment (i.e., impairments in social-communication and interaction, alongside repetitive stereotyped behavior and sensory anomalies)(1). Autism is well recognized as being highly heterogeneous on multiple levels: for example, in terms of its clinical presentation and underlying neurobiology. Indeed, more than 100 genes(2) and many aspects of brain structure have been associated with ASD at the group level(3). Autism is also grounded in the process of brain maturation and it is believed that alterations are evident throughout brain development(4, 5). In particular, differences in cortical thickness have been reported across different studies and ages(6), which – together with differences in surface area(6–10) – underpin regional differences in brain volume in ASD(11–13). However, the precise etiology of the disorder in terms of brain development and underlying mechanisms remain elusive.

The heterogeneity of ASD is a fundamental barrier to understanding the neurobiology of ASD and the development of interventions(14). Regional group-level differences have been reported across several neuroanatomical measures, including cortical thickness (CT)(8, 10, 15–22). However these findings show generally poor replication across studies(3, 7, 19, 23, 24) and small effect sizes (8, 19). Heterogeneity is also evident in studies that have used classifiers to discriminate ASD participants from controls, which mostly show relatively low accuracy for predicting diagnosis, especially in large samples(19, 25, 26). An important reason for this is that most studies to date have employed a traditional case-control approach, which is based on the assumption that the clinical and control groups are homogeneous entities(7, 27). Thus, the case-control approach provides information about alterations at the group level or, in other words, in the ‘average ASD participant’. However, different participants may have different symptom profiles and different etiological pathways and resulting neurobiological changes may converge on the same symptoms. Therefore, to understand the neurobiology of ASD, it is important to understand the range of associated neurobiological variation, which may subsequently inform intervention at the level of the individual in the spirit of ‘precision medicine’(28). A common approach to study the biological heterogeneity underlying ASD is to find subtypes using clustering algorithms, mostly on the basis of symptoms or behavioral characteristics(29–34). This approach has been somewhat successful and is appropriate if the clinical cohort can be cleanly partitioned into a relatively small number of homogeneous subgroups on the basis of the chosen measures. However, it does not tackle heterogeneity within subgroups and it may be the case that no clearly defined subgroups exist in the data. Moreover, subgroups derived from behavior or symptoms require extensive validation on external measures and still may not fully reflect the underlying biology(35, 36).

Here, we apply a complementary normative modelling approach(36, 37) to understand the biological heterogeneity of ASD. This shifts the focus away from group-level comparisons – which can detect consistent differences across groups of individuals (e.g. diagnoses or putative subtypes) – towards characterizing the degree of alteration in each individual, with reference to the typically developing brain. This allows us to detect and map neuroanatomical alterations at the level of the individual and has recently shown promise in understanding the biological variation of psychotic disorders(37). Normative modelling is analogous to the use of growth charts in pediatric medicine, which allow the development (e.g. in terms of height or weight) of each individual child to be measured against expected centiles of variation in the population. To achieve this, we first estimated a statistical model characterizing typical cortical development that accurately quantifies the variation within the population and across brain development. We then placed each individual ASD participant in relation to the typical distribution in order to identify alterations in individual cases with respect to the typical pattern of brain maturation. Our main goals were to: (ii) to map the neuroanatomical features by which each individual ASD participant differs from the expected typically developing pattern, across both different developmental stages and levels of functioning and thereby (ii) demonstrate the value of normative modelling techniques for understanding the biological heterogeneity of ASD. For this, we employed data from a large international study(38) with harmonized data acquisition procedures and a design that naturally groups of subjects according to different developmental stages. Whilst normative modelling is suitable for many different aspects of brain structure or function, here, we focused on cortical thickness (CT) which is a sensitive and reliable measure of cortical morphology in ASD(6, 8, 39), although we also investigated surface area (SA). Ultimately, we hope this approach will yield a set of individualized neurobiological ‘fingerprints’ facilitating a route towards precision medicine approaches in ASD(28).

## Materials and Methods

### Participants

Full details on study design and clinical characteristics have been described previously(38). Briefly, we included all participants from the Longitudinal European Autism Project (LEAP)(40) cohort with a structural MRI scan surviving quality control and the necessary clinical and demographic data. We included 206 typically developing (TD) individuals aged 7 to31 years (127 male; Table 1; Table S1, Figure S1) and 321 individuals aged 6 to31 years (232 male) with ASD. There were no significant differences between TD and ASD cohorts in age but the IQ of ASD participants was lower than TD participants. Under the study design, each cohort was split into four subgroups according to age and level of intellectual ability (Table 1): (i) Adults with ASD without intellectual disability (ID) and TD controls aged 18 to 30 years, IQ ≥ 70; (ii) Adolescents with ASD without ID and TD aged 12 to 17 years; (iii) Children with ASD without ID or TD aged 6 to 11 years; and (iv) Adolescents and adults with ASD and ID (i.e. full-scale IQ between 50 to 70(1)) aged 12 to 30 years. Note that only TD participants were included in the estimation of the normative model.

**Table 1.**
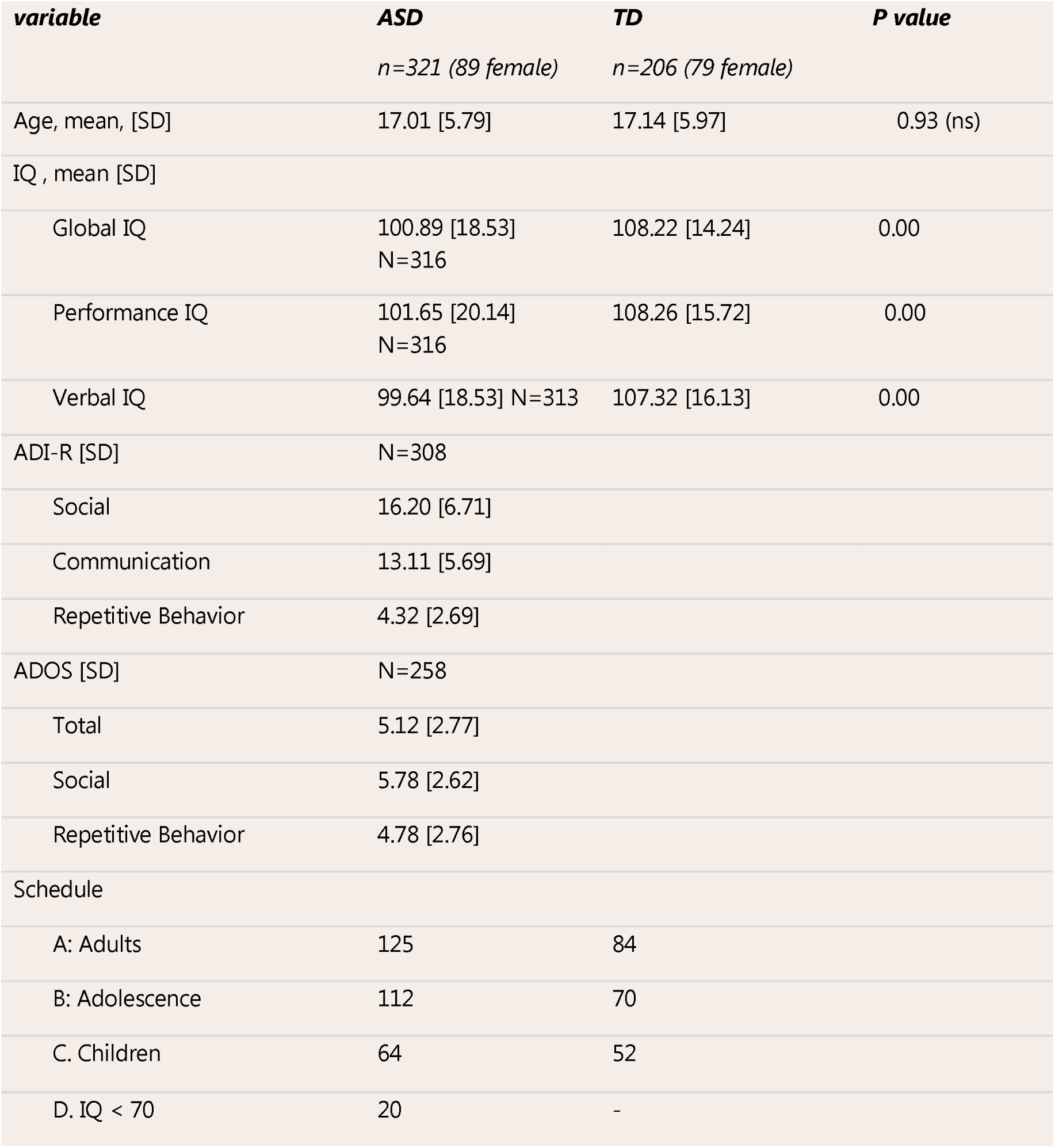
Clinical characteristics

| **variable** | **ASD**<br>*n=321 (89 female)* | **TD**<br>*n=206 (79 female)* | **P value** |
| --- | --- | --- | --- |
| Age, mean, [SD] | 17.01 [5.79] | 17.14 [5.97] | 0.93 (ns) |
| IQ , mean [SD] |  |  |  |
| Global IQ | 100.89 [18.53]<br>N=316 | 108.22 [14.24] | 0.00 |
| Performance IQ | 101.65 [20.14]<br>N=316 | 108.26 [15.72] | 0.00 |
| Verbal IQ | 99.64 [18.53] N=313 | 107.32 [16.13] | 0.00 |
| ADI-R [SD] | N=308 |  |  |
| Social | 16.20 [6.71] |  |  |
| Communication | 13.11 [5.69] |  |  |
| Repetitive Behavior | 4.32 [2.69] |  |  |
| ADOS [SD] | N=258 |  |  |
| Total | 5.12 [2.77] |  |  |
| Social | 5.78 [2.62] |  |  |
| Repetitive Behavior | 4.78 [2.76] |  |  |
| Schedule |  |  |  |
| A: Adults | 125 | 84 |  |
| B: Adolescence | 112 | 70 |  |
| C. Children | 64 | 52 |  |
| D. IQ < 70 | 20 | - |  |

TD participants were recruited via advertisement. Individuals with an existing ASD and/or mild ID diagnosis (according to DSM5/ICD10 criteria) were recruited from existing databases and clinic contacts across one of seven study sites: the Institute of Psychiatry, Psychology and Neuroscience, King’s College London, UK, Autism Research Centre at the University of Cambridge, UK, Radboud University Nijmegen Medical Centre, University Medical Centre Utrecht, The Netherlands, Central Institute of Mental Health, Mannheim, Germany, and the University Campus Bio-Medico, Rome, Italy. The combined information of Autism Diagnostic Interview-Revised(41) (ADI-R) and Autism Diagnostic Observation Schedule Second Edition(42) (ADOS-2) were used to measure symptom severity(33). However, individuals with a clinical ASD diagnosis who did not reach conventional cut-offs on these instruments were not excluded. The ADI-R is a parent reported measure of lifetime or past developmental window symptom severity whereas the ADOS-2 is an expert rating of current symptoms. A standard set of exclusion criteria were applied and are provided in the supplementary material. All subjects were scanned with a T1-weighted imaging protocol and Freesurfer (version 5.3) was used to estimate measures of regional cortical thickness and surface area. See supplementary methods for details.

### Constructing a normative model of cortical thickness

An overview of the normative modelling approach is shown in Figure 1 and has been described previously(36). Briefly, Gaussian process regression (GPR)(43) was used to estimate separate normative models of CT and SA at each vertex on the cortical surface (see supplementary methods for details). This normative model can be used to predict both the expected cortical thickness and the associated predictive uncertainty for each individual participant. The contours of predictive uncertainty can then be used to model centiles of variation within the cohort. This allows us to place each individual participant within the normative distribution thereby quantifying the vertex-wise deviation of cortical thickness from the healthy range across the brain.

**Figure 1:**
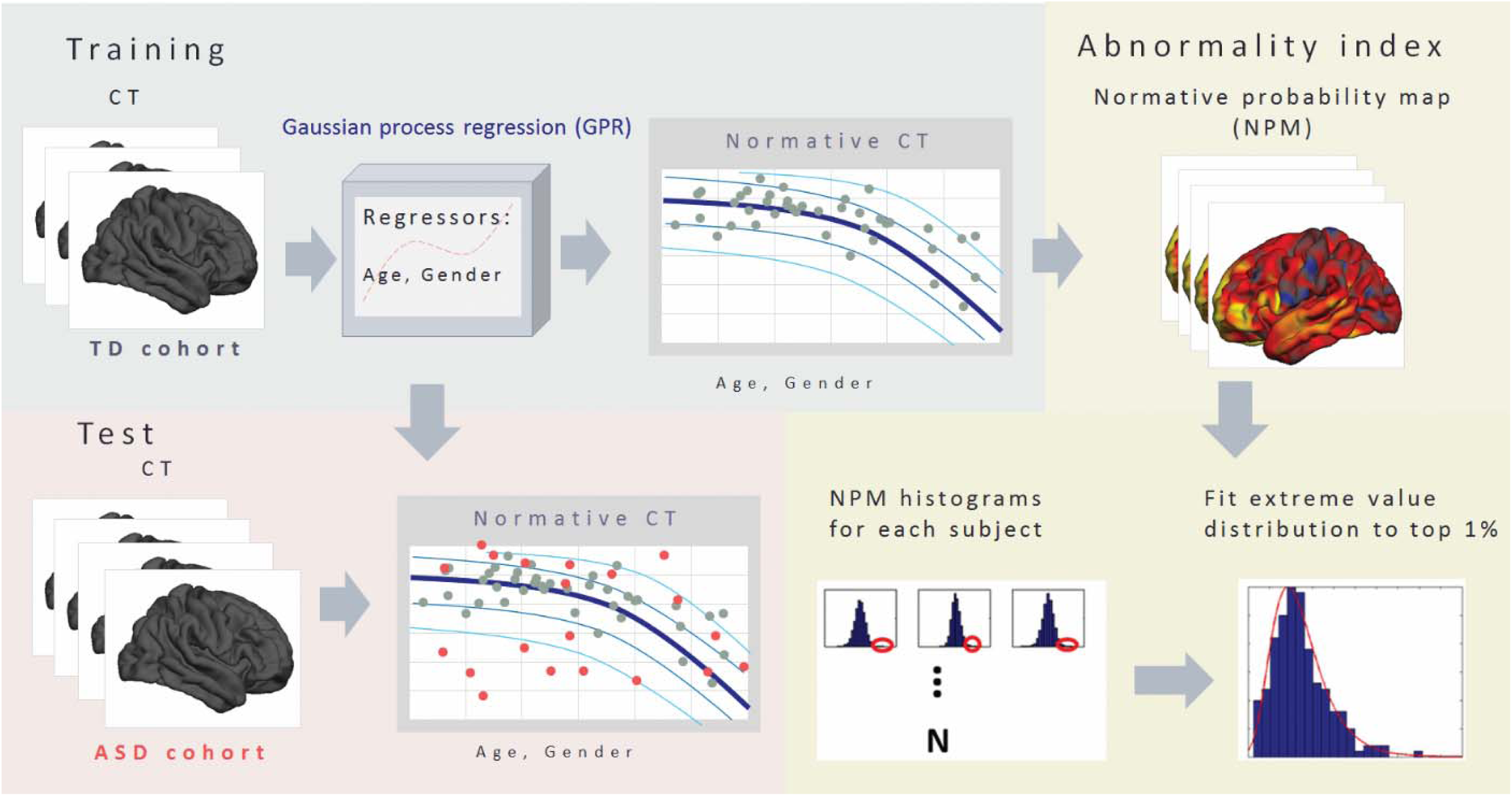
Methodological overview. First, a normative model was estimated from cortical thickness derived from TD subjects (gray dots). Then we use this model to predict cortical thickness in ASD subjects (red dots). This allows us to estimate normative probability maps which show the regional deviations from the expected pattern in each subject. Finally, we generate a summary statistic quantifying the overall deviation for each subject by taking maximum deviation across brain using extreme value statistics. Abbreviations: CT = cortical thickness, NPM = normative probability map. See text for details.

To achieve this, we generated a developmental model of typical brain development by training a GPR model on the TD cohort (N=206) using age and gender as covariates (i.e. independent variables) to predict cortical thickness (i.e. dependent variable). In pediatric medicine, growth charts are normally estimated on the basis of a large population cohort (i.e. potentially including patients with various disorders based on the population prevalence). In our sample, the prevalence of ASD is much higher than in the population, so for simplicity and to avoid the normative model being enriched for ASD, we estimated the normative model on the basis of the TD participants only. Moreover, while the number of data we employ here is relatively small in comparison with population based studies, our Bayesian statistical model provides a principled method to handle uncertainty and therefore automatically makes inferences more conservative as the number of data points decreases, although more data would allow more precise estimates. To assess generalization, we used 10-fold cross-validation before retraining the model using the whole dataset to make predictions on the ASD participants following standard practice in machine learning (see supplementary methods for details). Importantly, all parameters were estimated using the training data using empirical Bayesian estimation (36) and the use of cross-validation ensures unbiased estimates for the TD cohort as well as for the ASD cohort. Therefore, deviations can be compared with one another.

### Estimating regional deviations for each subject

To estimate a pattern of regional deviations from typical cortical thickness for each participant, we derived a normative probability map (NPM) that quantifies the deviation from the normative model for cortical thickness at each vertex. This was done by using the normative model to predict vertex-wise estimates of cortical thickness for each individual participant, then estimating a subject-specific Z-score (36) (supplementary methods). This provides a statistical estimate of how much each individual differs from the healthy pattern at each vertex. We thresholded the NPMs, correcting for multiple comparisons by controlling false discovery rate (FDR) at p < 0.05 within each participant, as in (36).

To measure the spatial overlap of the individualized deviations across the cohort, we calculated an overlap map by counting the significant (FDR corrected) vertices derived from the Z-score maps across all subject-level NPMs. The resulting summary maps indicate the spread of vertex-wise deviations across the brain, separately for positive and negative deviations. This allowed us to identify a set of brain regions where participants had increased (positive deviation) or decreased (negative deviation) cortical thickness relative to the reference cohort.

To provide a simple comparison for these subject-level deviations, we also estimated a standard vertex-wise general linear model to establish significant differences between groups including age as a covariate. We also investigated models including quadratic and cubic age terms (corrected using FDR at p < 0.05) and separate models for males and females.

### Constructing an individual-level atypicality score

A key benefit of normative modelling is a probabilistic interpretation of the deviations across all subjects. The NPMs therefore provide a multivariate measure of deviation from the normative range across all brain regions. This captures spatially distributed differences from the TD pattern. To better understand most important focal differences for each subject we estimated a summary score for each participant capturing the individual’s largest deviation from the typical pattern (which is potentially the most clinically relevant). This can be modelled using extreme value statistics (44) and is based on the notion that the expected maximum of any random variable converges to an extreme value distribution (EVD). Therefore, we estimated a maximum deviation for each subject by taking a trimmed mean of 1% of the top absolute deviations for each subject across all vertices and fit an EVD to these deviations.

### Mapping behavioral associations

Last, to assess the clinical relevance of these deviations, we computed Spearman correlation coefficients between global and regional extreme deviation from the normative model and ADOS-2/ADI-R symptom severity scores (p < 0.05, FDR). The global measure (described above) provides an overall summary of the deviation for each individual whilst the regional assessment helps to determine the functional correspondence of the deviations across individuals on a region-by-region basis. The regional extreme deviation was computed as the trimmed mean of the 1% of top absolute deviations for each region after parcellating the cortex using the Desikan-Killiany atlas(45).

### Checking for potential confounds

To investigate whether potential confounds could have influenced our findings, we estimated a separate normative model additionally including dummy regressors for IQ and site. We also performed post-hoc tests between the deviations from the normative model and potential confounding variables (IQ, comorbid symptoms, and surrogate measures of image quality) See supplementary Tables 2 and 3.

## Results

### A normative model quantifying the decline of cortical thickness with age

Figure 2 shows the developmental normative model of CT derived from the TD male cohort, thresholded to show vertices where the correlation between true and predicted labels was higher than predicted by chance (p < 0.05, FDR corrected. See Figure S3 for females). The unthresholded map showing the correlation between true and predicted CT values is shown in Supplementary Figure S1 along with the root mean squared error of the normative model across different vertices. In most regions, CT decreases consistently and approximately linearly with age. However, in some regions, cortical thickness followed a non-linear (i.e. an inverted U-shaped) trajectory with an early rise followed by a decline, e.g. in the inferior temporal and posterior frontal regions. This corresponds well with the known developmental trajectory of cortical thickness(46–50). The normative model for SA showed a similar, relatively global pattern of decline as for CT (not shown).

**Figure 2:**
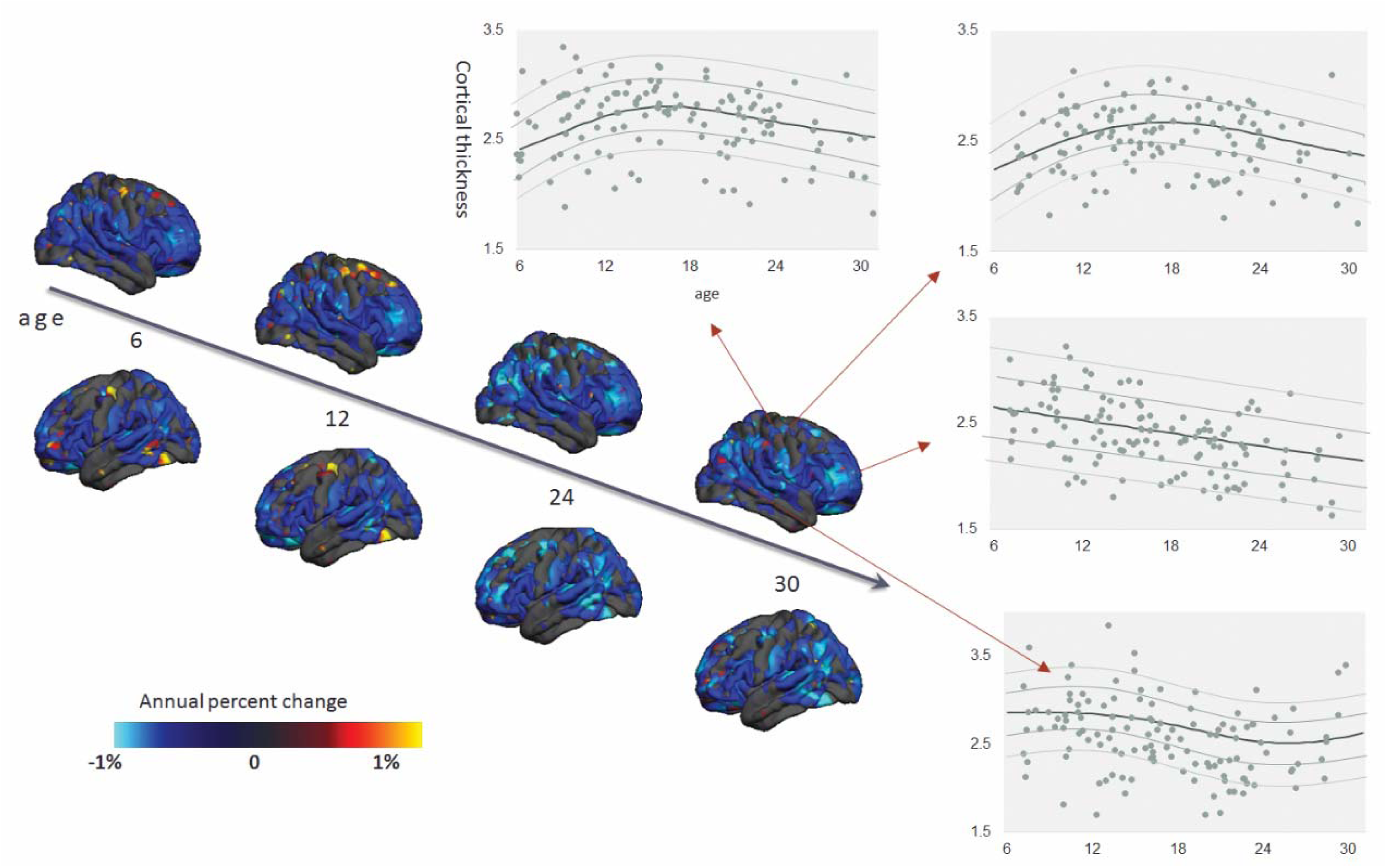
Normative model of developmental changes of cortical thickness across the developmental range in the typical developing male cohort (the model was estimated using both genders). Cortical thickness was predicted using a trained normative model across the age range of six to thirty-one. The predicted cortical thickness map was thresholded so that only vertices that could accurately predict the true cortical thickness in the healthy cohort under cross-validation were retained (Pearson correlation, p < 0.05, FDR). Blue vertices and yellow indicate reduced and increased CT respectively. Moreover, the predicted cross-sectional developmental trajectories of CT in four randomly-selected vertices are shown.

### Widespread deviations from normative pattern of cortical thickness among the ASD cohort

Figure 3 shows the classical mass-univariate group difference (i.e. case control) map between ASD and TD cohorts. This shows few significant differences between groups; only two small regions of increased CT in superior frontal and parietal cortices survived FDR correction. There were also few significant differences when additionally including quadratic and cubic age terms and no differences in the age x diagnosis interaction. The separate models for males and females also did not show any significant differences after FDR correction.

**Figure 3:**
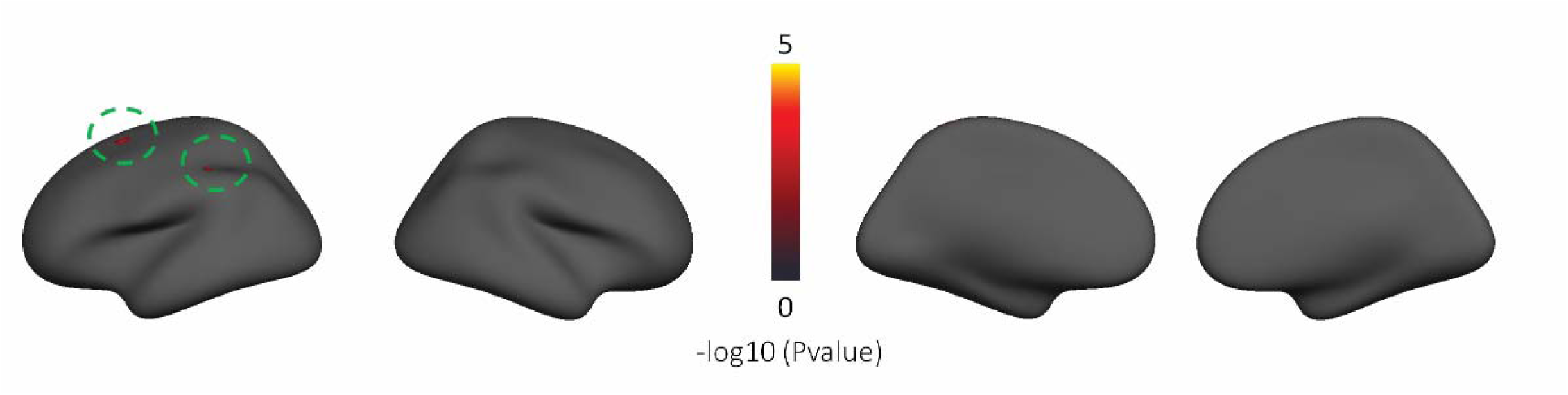
Vertex-wise group differences between ASD and TD cohorts after FDR correction (p < 0.05). The green circles indicate the regions show the vertex-wise group difference. No vertices survived after FDR correction in vertex-wise group differences map between ASD and TD in female and male separately.

Figure 4 and Figure 5 show a summary of the NPMs for the ASD and TD cohort. Specifically, these figures show the number of participants in each group that deviate negatively (Figure 4) or positively (Figure 5) from the normative model at each vertex after intra-individual FDR correction. Importantly, and contrast to the GLM, these deviations need not overlap between subjects. As expected, the TD cohort shows few significant deviations, indicating that the normative model provides a good fit for this cohort. Crucially, this fit was achieved under cross-validation and is therefore unbiased. Therefore, under the ‘null’ hypothesis that ASD participants follow a similar trajectory of brain development to TD participants there is no prior reason to expect the fit will be better in TD than in ASD participants. In contrast, the total number of deviating vertices was noticeably higher in the ASD cohort and was widespread across the brain, suggesting that there are widespread and individualized deviations from the normative model in certain subsets of participants. When considering each age group separately, negative deviations were most prominent in children, whereas positive deviations were most prominent in adolescents and adults. The results were very similar for the models including IQ and scanning site as covariates (supplementary Figures S4 and S5) and a similar pattern of results was observed for SA, albeit with slight differences with respect to the pattern of deviations across brain regions (supplementary Figure S6 and S7).

**Figure 4:**
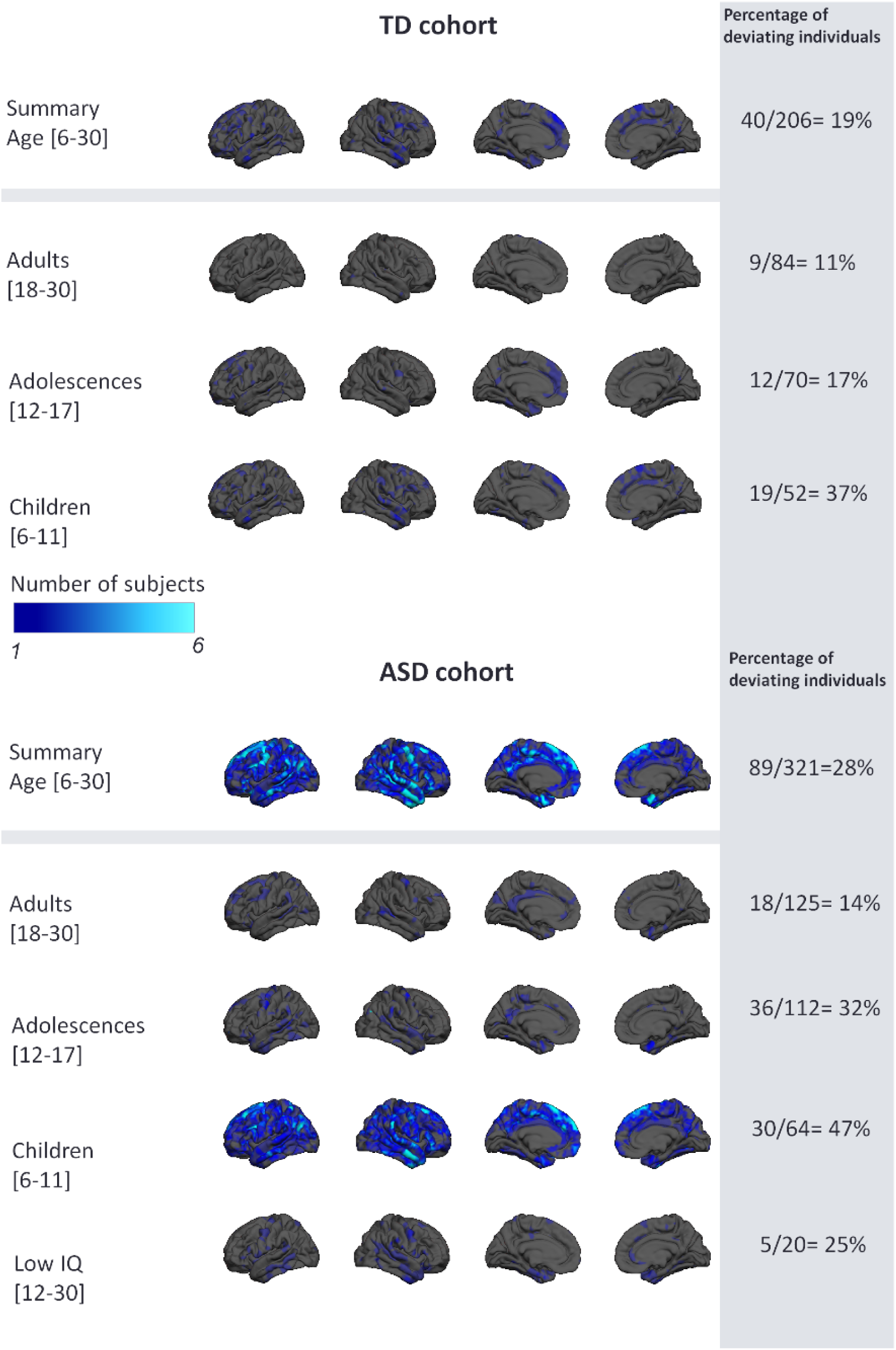
Overlap of vertex-wise negative deviation across each cohort and schedule. This map shows the spatial distribution of individual subjects with significant deviations in each vertex after FDR correction. The proportion of subjects contributing to each map is also shown (i.e. the proportion of subjects having deviations surviving FDR correction).

**Figure 5:**
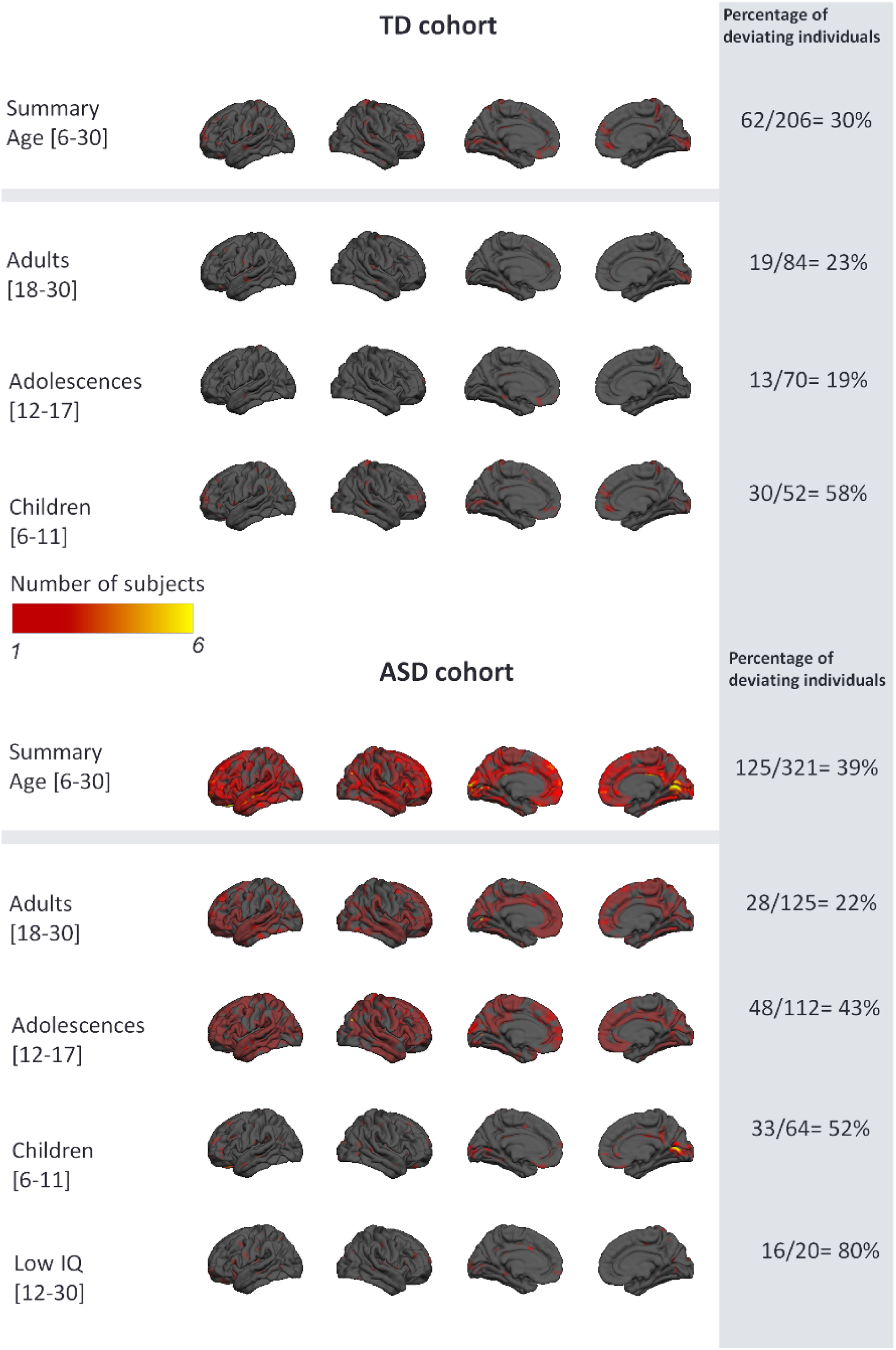
Overlap of vertex wise positive deviation across each cohort and schedule. See caption to Figure 4 for further details

### ASD participants deviate more than TD from the normative pattern of development

Figure 6 shows the distribution of the most extreme deviations from the normative model across the brain. This shows that the maximum deviation across the brain is higher in the ASD cohort relative to the TD cohort and the distribution of the ASD cohort is shifted towards the right implying relatively more subjects with extreme deviations. Saliently, the top fifteen deviating individuals belong to the ASD cohort, which is extremely unlikely to occur by chance (p <0.0005, binomial test). The NPMs of these participants (supplementary Figure S8) have highly individualized patterns of deviation not only with respect to brain regions but also in sign, with some participants having positive deviations (i.e., greater CT) or negative deviations (reduced CT). These participants did not show a consistent pattern with respect to their symptom scores (supplementary Table 5), which underscores the degree of clinical and neurobiological heterogeneity within the ASD cohort. However, with regard to their demographic profile, subjects with predominantly positive deviations were adolescents or adults, whilst most subjects with negative deviations were children.

**Figure 6:**
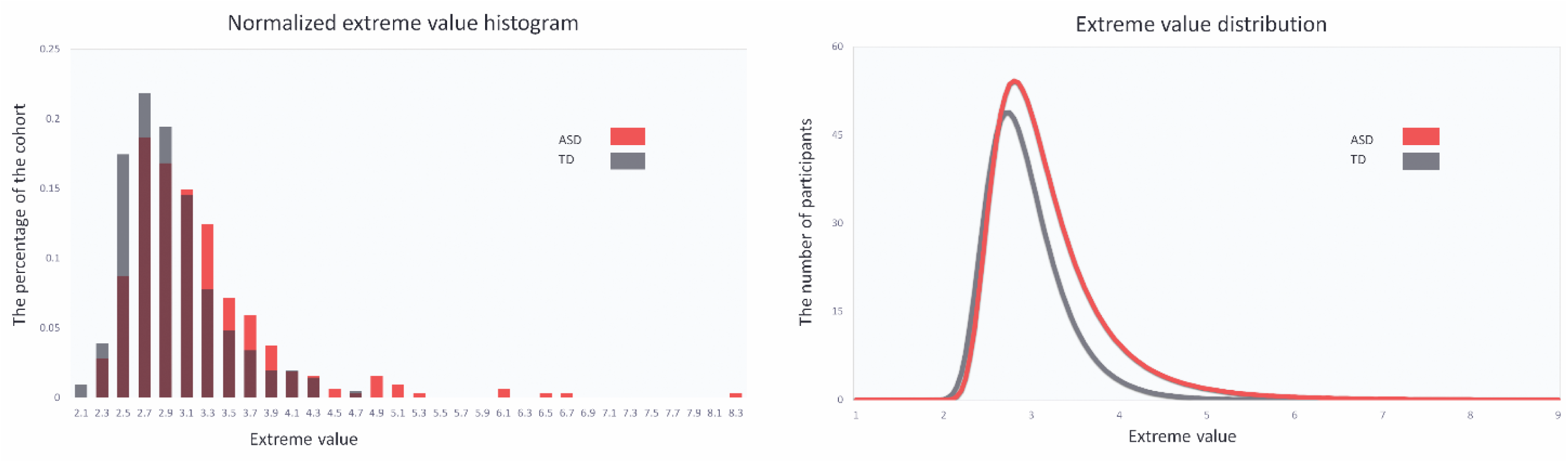
Extreme value histogram and distribution

### Association with symptoms

Global deviations from the normative model were negatively associated with ADOS repetitive behaviours (ρ=-0.21 p < 0.05) and regional deviations were associated with symptoms in several brain regions (Figure 7 and Figure 8). Associations were found with symptom severity in the repetitive domain of the ADOS-2 or ADI-R in prefrontal regions in females. In males, a similar pattern was seen, but did not survive multiple comparison correction except the superiorfrontal region in ADI-R. Social interaction and communication scores also had nominally significant associations in females but these did not survive correction.

**Figure 7:**
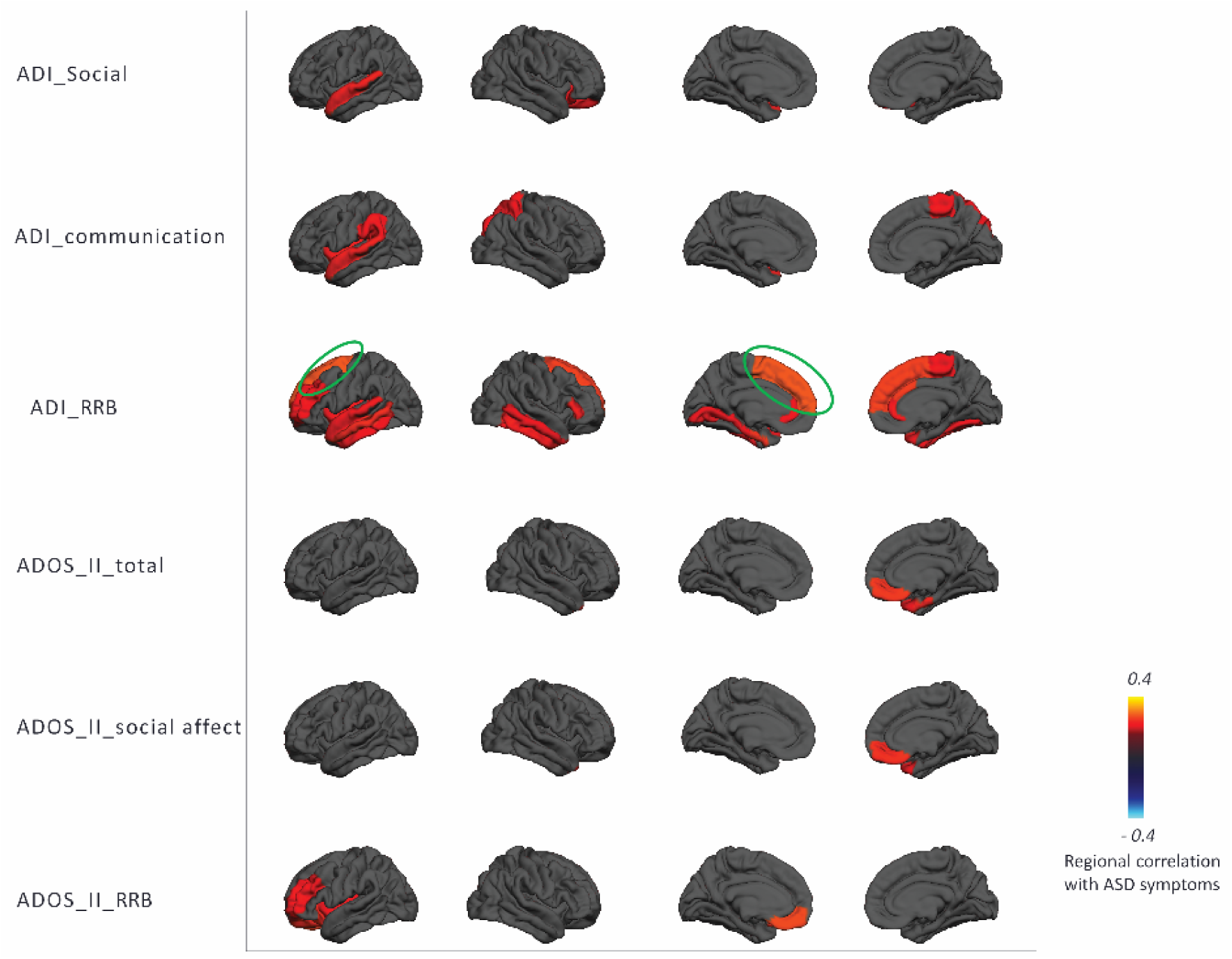
Regional extreme value deviation correlation with ASD symptoms-female (p <0.05) according to the Desikan-Killany parcellation scheme. Blue and yellow regions indicate negative and positive association with ASD symptoms, respectively. Green circles indicate the regions that survived after FDR correction.

**Figure 8:**
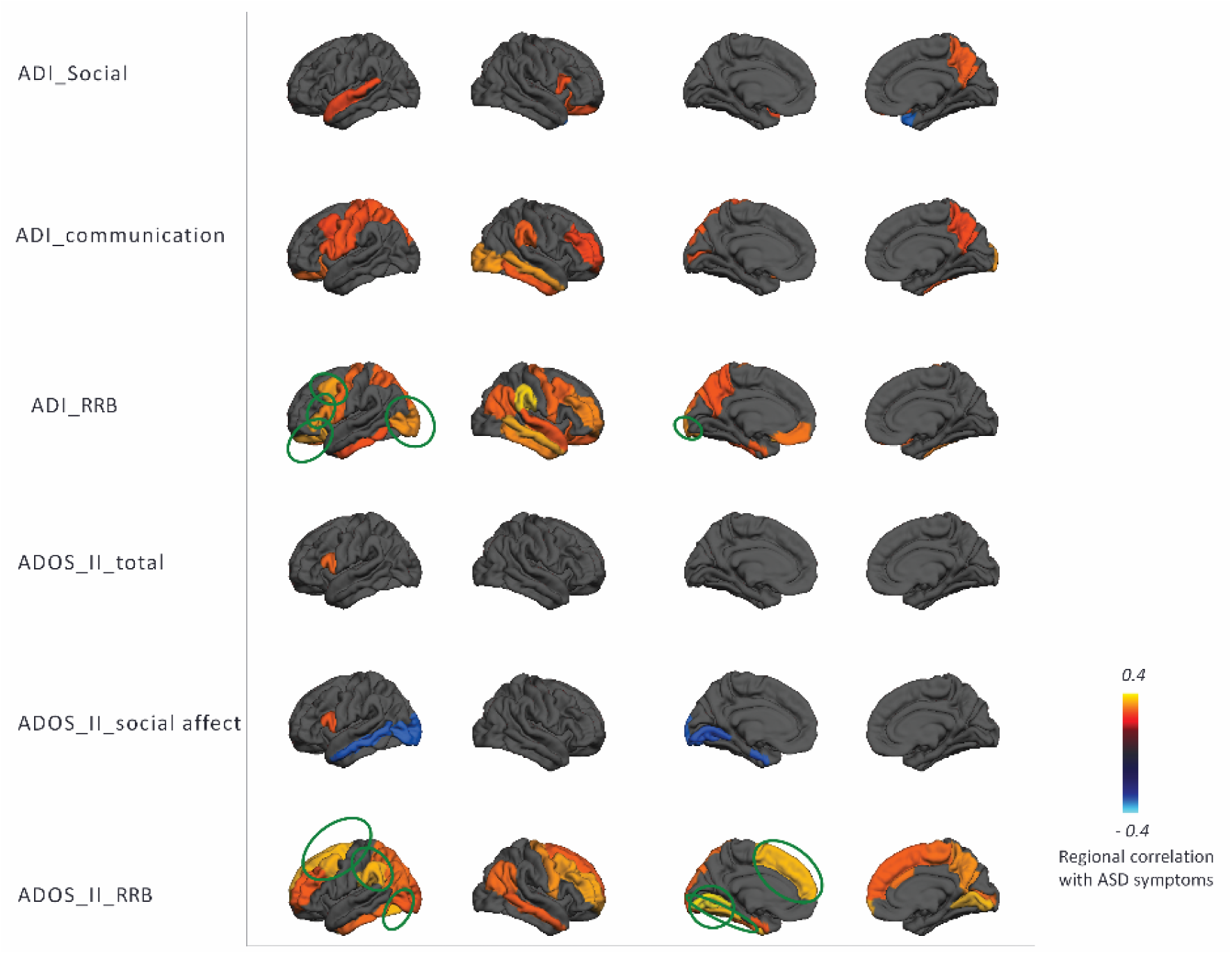
Regional extreme value deviation correlation with ASD symptom for males (p <0.05) according to the Desikan-KIllany parcellation scheme

## Discussion

In this study, we aimed to dissect the heterogeneous neurobiology of ASD by mapping the deviation of each individual participant from a normative model of cortical thickness development. In a large heterogeneous cohort spanning a wide range of the ASD phenotype, we showed few significant group-level differences between ASD and TD cohorts in cortical thickness using a classical case-control analysis. In contrast, our normative modelling approach showed striking, widespread patterns of cortical atypicality at the level of individual ASD participant. These patterns were highly individualized across participants, distinct across different developmental stages and associated with symptoms, especially repetitive behaviors. This supports the notion that a subset of ASD participants follow a different developmental trajectory to TD subjects, and that the trajectory each ASD participant follows is highly individualized. From a methodological standpoint, our study shows that: (i) that it is necessary to look beyond the case control paradigm to understand the heterogeneous neuroanatomy of ASD; (ii) that normative modelling provides as an alternative conceptual framework for understanding the heterogeneous neurobiology of ASD in terms of deviations from a typical pattern and (iii) that focusing on an ‘average autistic individual’ provides only a partial reflection of the nature of the condition. In other words, the case control approach focuses on common effects rather than inter-individual variation. Capturing and capitalizing on such variation at the individual level is at the heart of precision medicine.

The normative model describes the variation in typical brain development showed a largely monotonic –and in some areas non-linear– decrease of cortical thickness throughout development, consistent with previous neuroimaging studies(46–49, 51–55). The fact that we observed widespread inter-individual differences between ASD participants in terms of their deviations from the normative model explains why our classical case-control analysis revealed few significant differences and why several large previous neuroimaging studies have also only detected relatively modest group level effects(8, 19). The heterogeneity underlying ASD is widely recognized(2, 56–62); some studies have reported reductions in cortical thickness in ASD(15) whereas some studies have reported increases.(16, 63). Saliently, these inconsistencies remain evident even in large studies; for example, a large study derived from the ENIGMA consortium demonstrated both regional increases and decreases in ASD at the group level that were consistent across development(8). Other studies – many derived from the ABIDE dataset(64) – have shown widespread increases in CT early in development that are attenuated later in development(19, 20, 47). Our results compliment these studies because of our focus on studying individual variation within the ASD cohort. We show that: (i) a subset of participants show decreased CT and SA in childhood whilst (ii) other patients show regional increases in childhood in different areas (e.g. peri-calcarine cortex); (iii) some of participants show increased CT and SA in adolescence or adulthood. Crucially however, these effects show minimal overlap across brain regions in different individuals. This is in line with another recent study applying normative modelling to ASD, which found effects in a subset of participants that were different from the main group effects (65). Thus, we consider that group-level effects can be understood as the background upon which individual variation is superimposed. The individualized deviations we report were mostly located in areas previously associated with ASD, such as the medial cortex including cingulate and dorsomedial prefrontal regions, lateral prefrontal and parietal cortices, temporal cortices and the hippocampal formation(6, 7, 63, 66, 67). Whilst some of these regions have been associated with social processing, the individual deviations in these regions were not associated with social interaction or communication symptoms at the group level. This could be for several reasons, for example the anatomical patterns associated with these symptoms may be expressed in other measures of cortical anatomy (e.g.(68, 69)) or in subcortical regions. Adults and adolescents had relatively fewer deviations, but these were positive (relatively increased CT and SA) and widespread across prefrontal and temporal cortices. Notably, we detected relatively few deviations in ASD with ID, which is important to exclude the possibility that these subjects were driving the effects described above. However, the ASD with ID group was relatively small (N = 20) so we do not draw strong conclusions about potential differences between ASD with- and without ID.

The 15 subjects with the most atypical anatomy all had ASD, which is extremely unlikely to occur by chance. Moreover, these participants had individualized brain alterations and clinical characteristics. At the group level, the regional deviations we detected from the normative model were associated with the severity of lifetime and current autistic symptoms (ADI-R and ADOS-2 respectively), demonstrating that our model predictions may be clinically relevant. The deviation from the normative range was most informative about repetitive behavior symptom severity in that the strongest correlations were between CT in prefrontal regions with restricted repetitive behaviors, especially in females and across both parental report via ADI-R and observer ratings of current symptoms via ADOS-2. These results broadly correspond with previous reports(6, 70, 71) and suggest that ASD may be more heterogeneous in males, but we are cautious about this interpretation because we did not test it directly. Taken together, our results add weight to the importance of considering ASD in the context of a model of typical brain development and at the individual level(39, 63, 67)

Our findings should be considered in the light of several limitations. First, the trajectories of brain development were based on cross-sectional data and should be validated in a longitudinal cohort. Longitudinal follow-up data are currently being acquired and will be the subject of a future report. Second, we registered all subjects to a standard adult template brain, as is standard in the field(10, 63, 67, 72–74), which could cause bias. However, there were few deviations in the TD cohort which makes this possibility unlikely. Third, our data does not permit strong inferences about the degree to which confounding variables may have influenced our findings. We found moderate associations between deviations from the normative models and a surrogate metric of image quality, but these were also associated with childhood ASD symptoms, comorbid ADHD symptoms and IQ. Moreover, our study design does not permit inferences about the direction of causality. For example, subjects with the most abnormal anatomy may also have the most impairment. Finally, we did not perform manual edits on the cortical surface reconstructions. Whilst this eliminates one potential source of bias, the results need to be interpreted in the light of this and it is possible that performing manual edits may improve the quality of the surface reconstructions in some cases.

In conclusion, we estimated a normative model of cortical development based on a large typically developing cohort and applied this model to a heterogeneous ASD cohort. Our results show that it is necessary to look beyond the case-control paradigm –which is limited to detecting group-level effects describing the ‘average ASD participant’– to understand the heterogeneous neurobiology of ASD. Normative modelling is well suited for this purpose as it can chart the individualized deviation of each individual subject relative to the normative range, and hence provides an excellent tool for understanding the heterogeneity of psychiatric disorders.

## Acknowledgements

We gratefully acknowledge the support of the EU-AIMS LEAP study team for data acquisition, quality control and preprocessed and also support from The Netherlands Organization for Scientific Research (NWO) through VIDI grants to AFM (Grant No. 016.156.415) and CFB (864.12.003). JKB received funding from the FP7 under Grant Nos. 602805 (AGGRESSOTYPE), 603016 (MATRICS), and 278948 (TACTICS) and from the European Community’s Horizon 2020 Programme (H2020/2014-2020) under Grant Nos. 643051 (MiND) and 642996 (BRAINVIEW). We also gratefully acknowledge funding from the Wellcome Trust UK Strategic Award (098369/Z/12/Z). This work was supported by EU-AIMS (European Autism Interventions), which receives support from the Innovative Medicines Initiative Joint Undertaking under grant agreement no. 115300, the resources of which are composed of financial contributions from the European Union’s Seventh Framework Programme (grant FP7/2007-2013), from the European Federation of Pharmaceutical Industries and Associations companies’ in-kind contributions.

