## Supplementary information for "Dissecting the heterogeneous cortical anatomy of autism spectrum disorder using normative models"

### Supplementary methods

#### Exclusion criteria

Exclusion criteria for the Longitudinal European Autism Project (LEAP) study included a history of substance abuse and standard neuroimaging contraindications (e.g. claustrophobia, metal implants). Individuals were also excluded if they had a history of bipolar disorder or psychosis. In contrast to case-control studies that aim to detect consistent group differences, here we were interested in characterizing the heterogeneity within autism spectrum disorder (ASD) at the level of the individual. Therefore, we did not exclude other comorbidities in the clinical group because up to 70% of ASD individuals have one or more psychiatric conditions(1) and 30-50% of individuals with ASD are on stable medications(2).

#### Magnetic Resonance Imaging

A high resolution T1-weighted image was acquired from each participant with a standard Alzheimer's disease neuroimaging initiative (ADNI) sequence (3), matched across scanning sites. Cortical thickness was estimated from the high-resolution T1-weighted image for each subject using Freesurfer version 5.3 (http://surfer.nmr.mgh.harvard.edu/). Prior to analysis, all surface reconstructions were visually assessed for reconstruction errors by at least 3 independent raters. We excluded a small number of scans with severe artifacts (e.g. caused by head motion). The rest of the scans were included 'as is', i.e. we did not allow manual edits to reduce the possibility of bias (e.g. due to individual differences in operator skill). Cortical thickness maps were then smoothed with a 10-mm surface-based Gaussian kernel.

#### Gaussian process regression

As mentioned in the main text, Gaussian process regression (GPR)(4) was used to estimate separate normative models of cortical thickness (CT) and surface area (SA) at each vertex on the cortical surface. Whilst other methods are also suited to this purpose (e.g. Bayesian polynomial regression), in preliminary testing we found that GPR provides superior estimation of the mean and the ability to map the variation across the cohort through centiles of predictive confidence. We refer the reader elsewhere for a full treatment of Gaussian processes(5, 6) but briefly, a Gaussian process (GP) specifies a distribution over functions, such that any finite number of elements has a joint Gaussian distribution. They are excellent tools for Bayesian regression: given a dataset specified by $\mathcal{D} = \{\mathbf{x}_i, y_i \in\}_{i=1}^{N}$ – where $\mathbf{x}_i$ are $D$-

dimensional vectors of covariates, $N$ is the total sample size and $y_i \in \mathbb{R}$ are response variables – the response variables are predicted using a potentially nonlinear regression model with additive Gaussian noise, i.e.: $y_i = f_i + \epsilon_i$ where $\epsilon_i \sim N(0, \sigma_n^2)$. Inference then proceeds by placing a GP prior over this function then computing the posterior distribution using the canonical GPR predictive equations(5). This prior is uniquely specified by a mean ($m(x)$) and covariance ($k(x, x')$) function. Here, without loss of generality we choose a mean function equal to zero and a generic covariance function combining linear and non-linear terms, i.e.:

$$k(\mathbf{x}_i, \mathbf{x}_j) = \mathbf{x}_i^T \mathbf{x}_j + \sigma_f \exp\left(-\frac{1}{2}(\mathbf{x}_i - \mathbf{x}_j)^T \mathbf{\Lambda} (\mathbf{x}_i - \mathbf{x}_j)\right)$$

Where $\sigma_f$ is a signal amplitude parameter for the nonlinear component and $\mathbf{\Lambda}$ is a diagonal matrix with $\ell_d^{-2}$ along the leading diagonal. These are 'automatic relevance determination' parameters(5) that can down-weight irrelevant dimensions in the input space or emphasize important dimensions. Training a GP model refers to finding the optimal values for the model parameters which are: $\ell_1, \dots, \ell_D, \sigma_n$ and $\sigma_f$. This is conveniently achieved by maximizing the logarithm of the model evidence (i.e. the denominator of Bayes rule). Finally, we compute a single subject Z-statistic image for each subject ($i$) and at each brain location ($j$) by computing:

$$z_{ij} = \frac{y_{ij} - \hat{y}_{ij}}{\sqrt{\sigma_{ij}^2 + \sigma_{nj}^2}}$$

Here, $\hat{y}_{ij}$ is the predicted mean and predicted variance, $\sigma_{ij}$, which is combined with the true response ($y_{ij}$) and variance learned from the TD distribution ($\sigma_{nj}$). Because we estimate a separate noise parameter for each vertex, this should accommodate regional differences in population variation (for example, the estimated variance parameter will be higher in the regions where there is greater variation across individuals.

### Cross-validation

To assess generalization, we used 10-fold cross-validation where we partitioned the data into 10 'folds' and repeatedly trained the model on 90% of the data, withholding the remaining 10% for estimating generalization performance. This was repeated 10 times so that each partition was excluded once. This procedure is standard in machine learning and is known to provide approximately unbiased estimates of the true generalization ability.

#### Post-hoc investigation of potential confounding variables

To investigate the potential confounding effect of various potential confounding variables, we performed several tests. As described in the main text, we first estimated a normative model for CT additionally including scanning site and IQ as covariates. In addition, we additionally performed several post-hoc tests for potential confounding variables including scan quality, IQ and comorbid attention deficit/hyperactivity disorder (ADHD) symptoms. However, we emphasize strongly that these should be considered as illustrative only, because our study design does not allow us to determine the direction of cause-effect relationships. For example, it is reasonable to expect that subjects that show the most atypical cortical anatomy may also express the highest level of symptoms, have the most intellectual impairment and be the most likely to suffer from comorbid symptoms. First, to assess the possibility of scan quality (e.g. due to excessive head motion in the scanner) influencing our results, we correlated (using Spearman correlation) the deviations from the model with the Freesurfer Euler number (EN)(7). The EN summarizes the topological complexity of the estimated cortical surface and has been proposed as a proxy measure of scan quality(8) . However, this is an indirect measure in that it does not model scan quality directly, it should be considered with caution since many other variables can potentially influence EN, including age, atypicalities in cortical anatomy and disorder severity. Therefore, we also correlated EN with age and with ASD symptoms. In addition, we correlated the deviations from the normative with measures of full-scale IQ (see (9) ) and with measures of comorbid ADHD symptoms derived from the Development and Well-being Assessment(10) . We refer the reader elsewhere for a detailed description of these measures (3, 9).

### Supplementary results

#### Age histogram

Figure S1 shows the distribution of subject ages across diagnoses.

*Figure 1: Histogram of age of Female and Male individuals  across TD and ASD cohort*

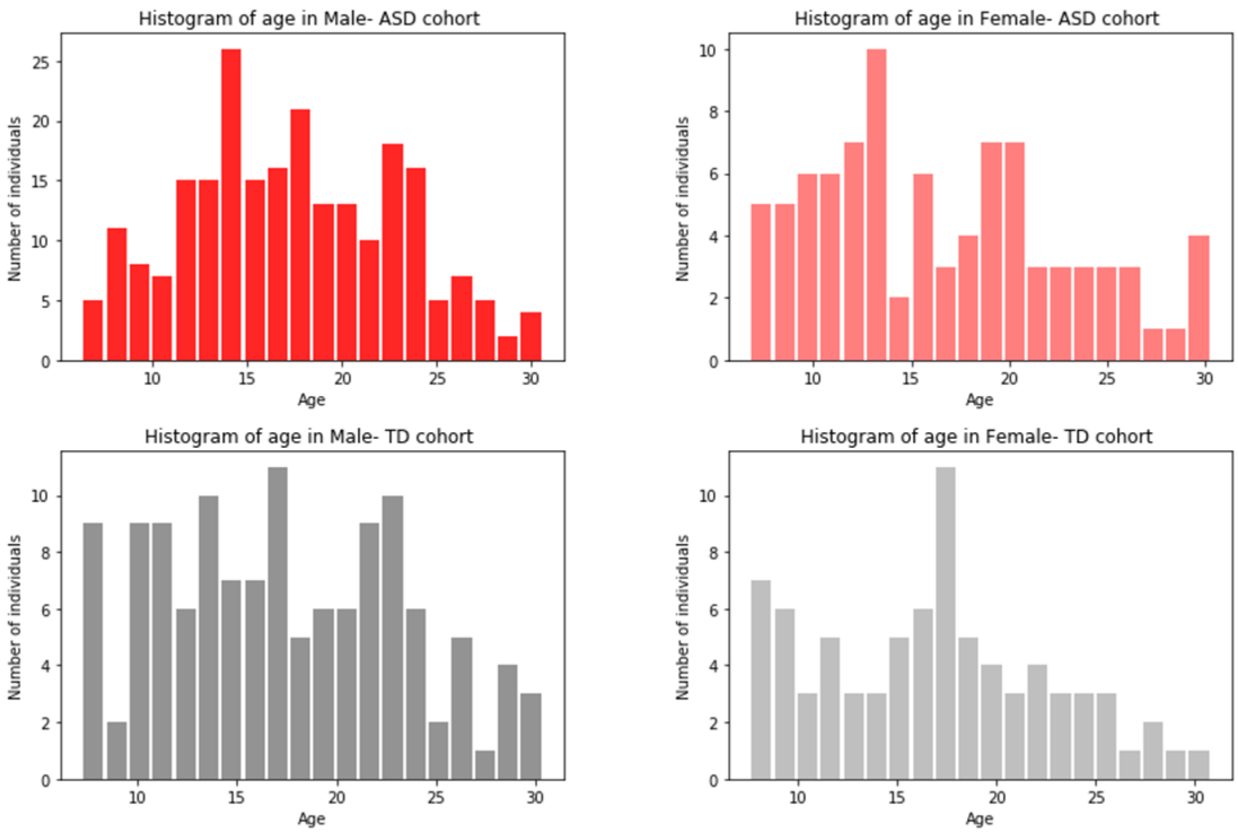

### Model fit evaluation

Figure 2 shows the mean accuracy of the normative model for predicting CT in typically developing (TD) and ASD participants, both in terms of root mean squared error (Figure 2A) and correlation between true and predicted CT values (Figure 2B).

*Supplementary Figure 2.A: Root mean square error of true and prective mean of cortical thickness in TD cohort*

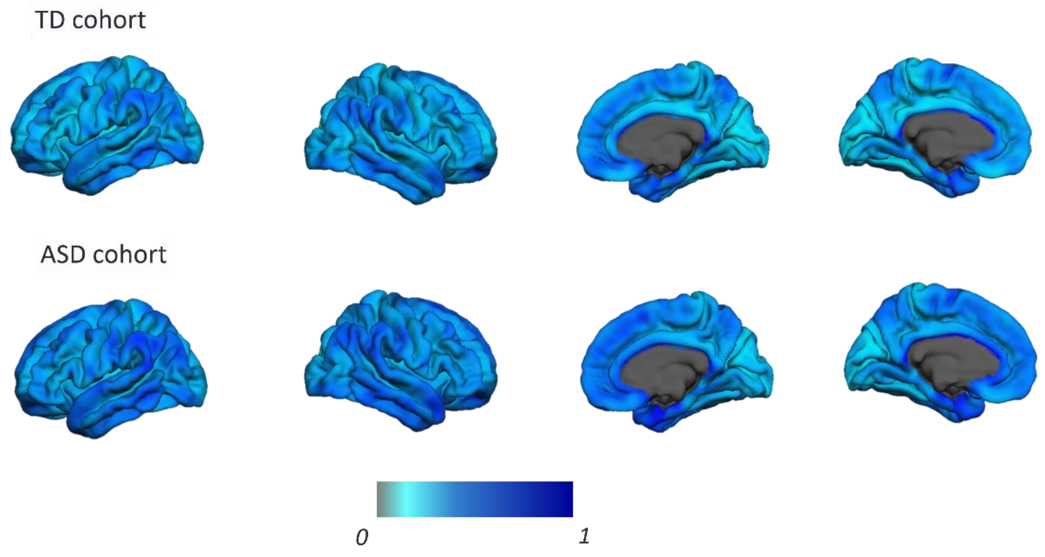


*Supplementary Figure 2.B: The correlation between true and prective mean of cortical thickness in TD and ASD cohort*

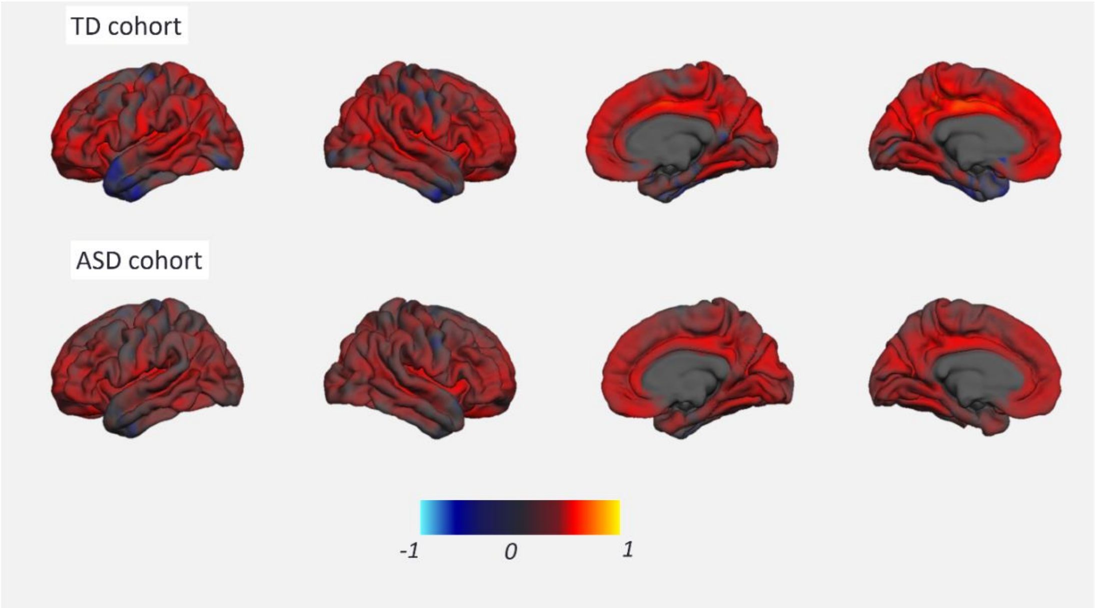

Normative developmental changes for cortical thickness in females

Supplementary Figure 3 shows the predictions made by the normative model for changes in female TD subjects (see Figure 2 in the main text for males).

*Supplementary Figure 3: Normative model of developmental changes of cortical thickness across the developmental range in the typical developing female cohort. Cortical thickness was predicted using a trained normative model across the age range of six to thirty-one. The predicted cortical thickness map was thresholded so that only vertices that could accurately predict the true cortical thickness in the healthy cohort under cross-validation were retained (Pearson correlation, p < 0.05, FDR). Blue vertices and yellow indicate reduced and increased CT respectively.*

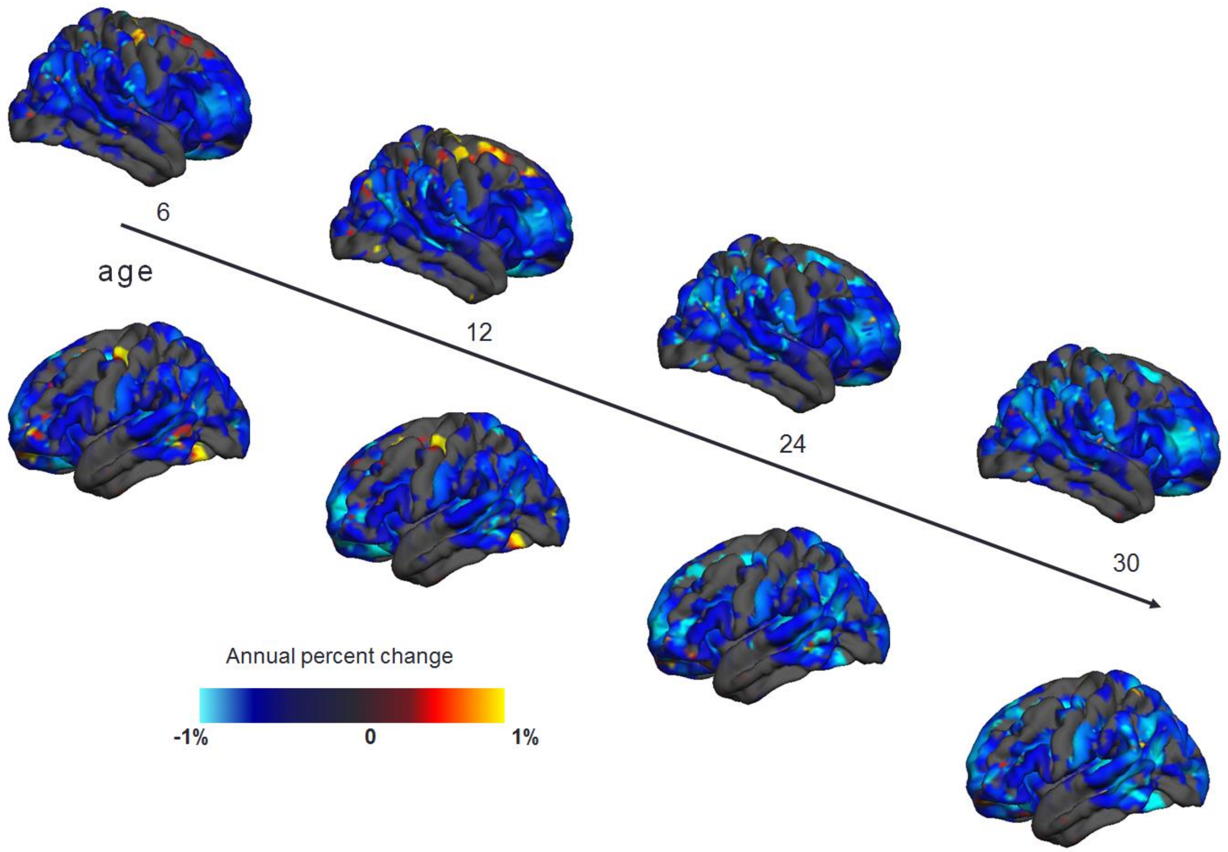

### Normative model of cortical thickness using age, gender, site and IQ as covariates

Supplementary Figures 4 and Figures 5 shows deviations from the normative model for CT re-estimated after additionally including IQ addition and scanning site dummy variables as covariates, separately for positive (Figure 4) and negative (Figure 5) deviations. The differences between this model and the original model are negligible and all the conclusions remain unchanged.

*Supplementary Figure 4: Overlap of vertex-wise negative deviation across each cohort and schedule. This map shows the number of subjects with significant deviations in each vertex after FDR correction*

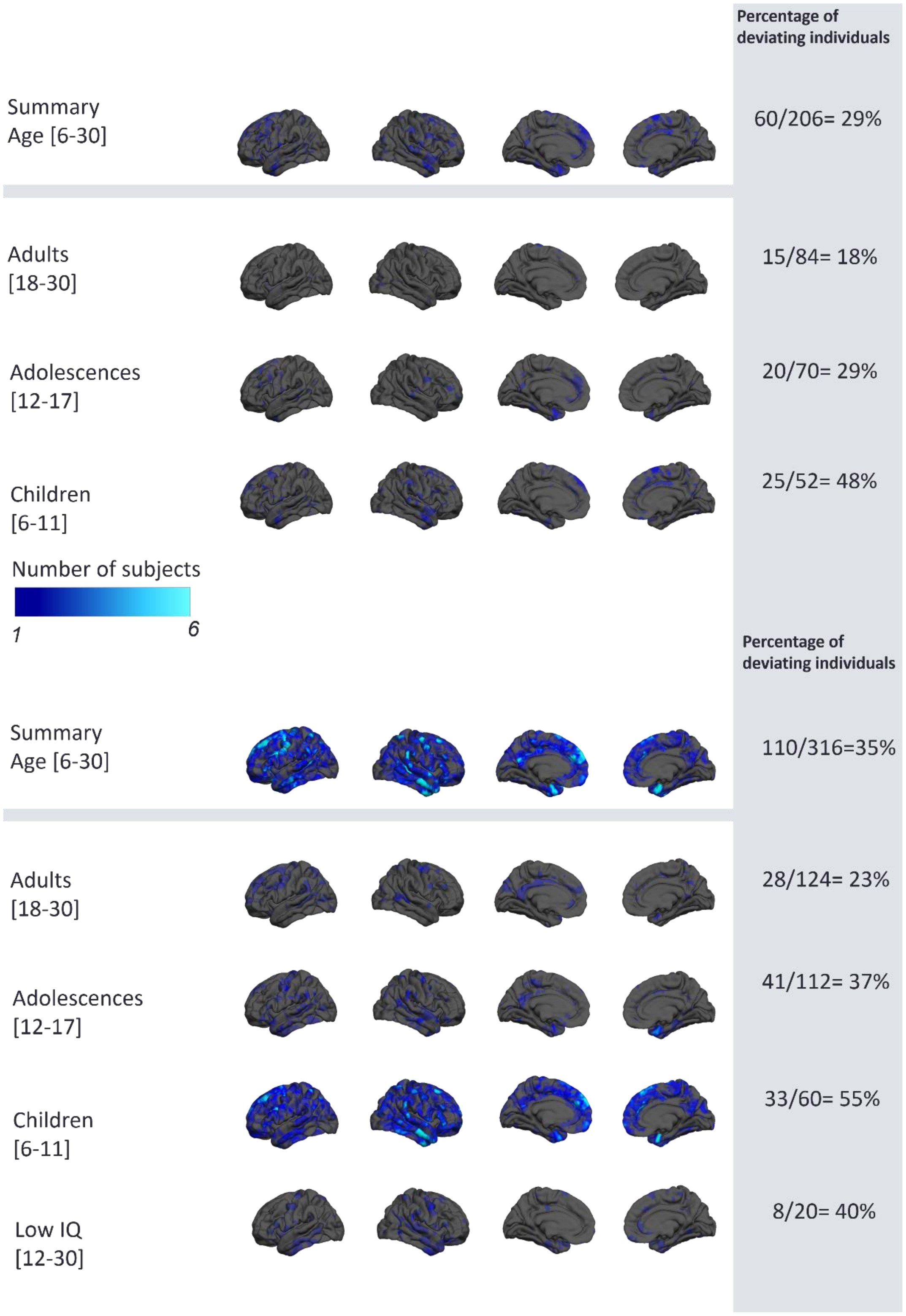

*Supplementary Figure 5: Overlap of vertex wise positive deviation across each cohort and schedule.*

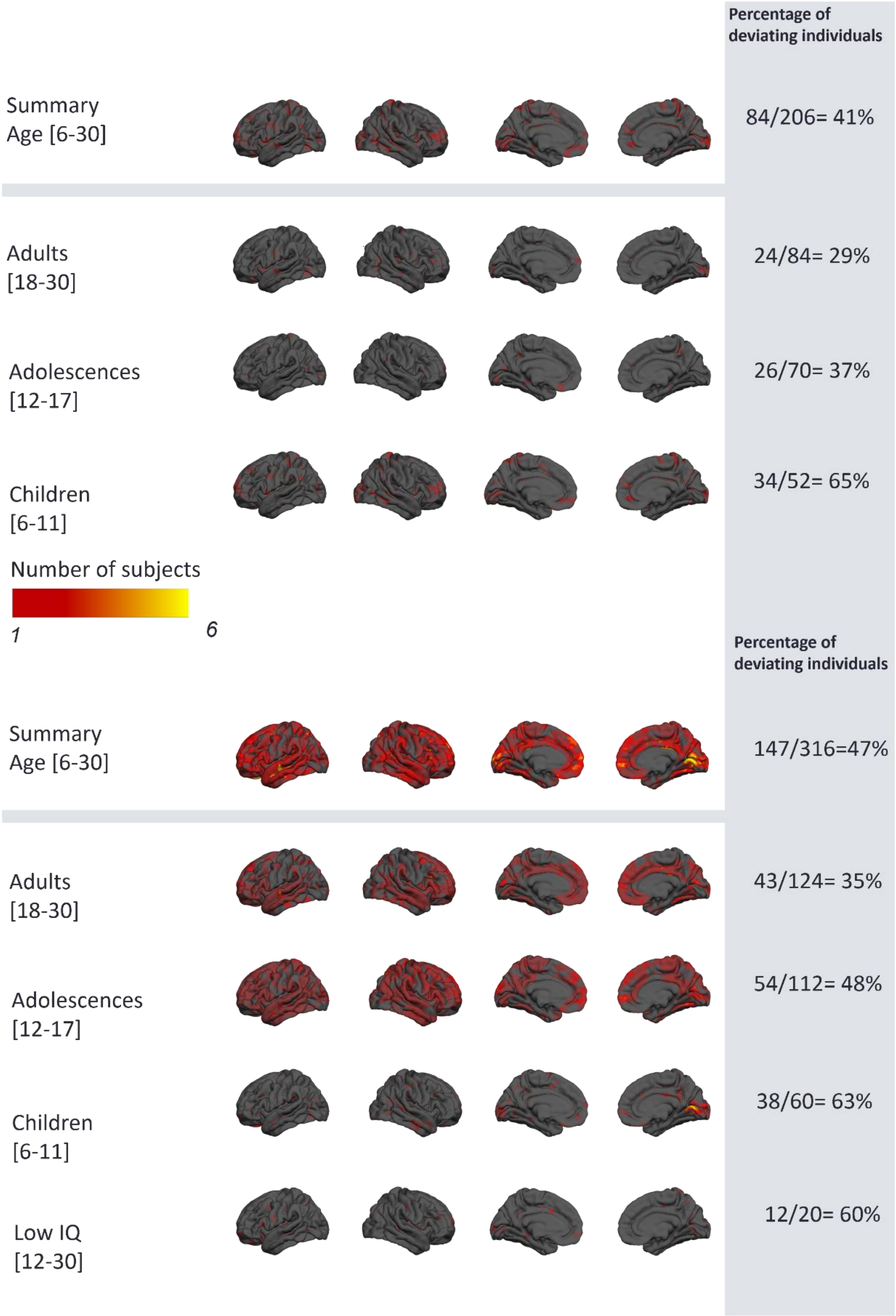

### Normative model of surface area

Supplementary Figure 6 and figure 7 shows deviations from the normative model for CT estimated using age and gender as covariates, separately for positive (Figure 6) and negative (Figure 7) deviations. The overlap of deviating voxels figures show a similar but slightly different pattern relative to CT.

*Supplementary Figure 6: Overlap of vertex-wise negative deviation in surface area normative model across each cohort and schedule. This map shows the number of subjects with significant deviations in each vertex after FDR correction*

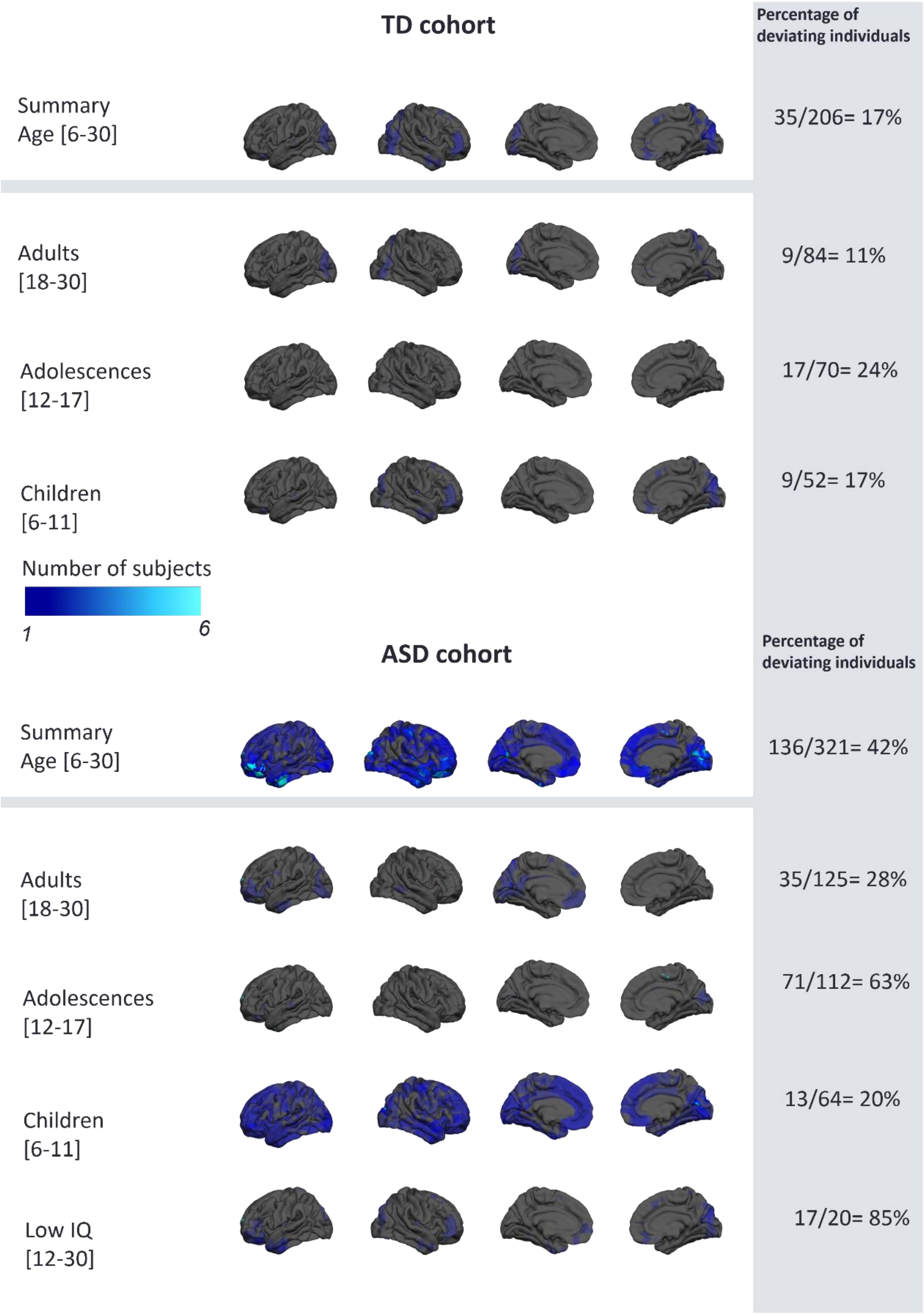

*Supplementary Figure 7:Overlap of vertex wise positive deviation across each cohort and schedule.*

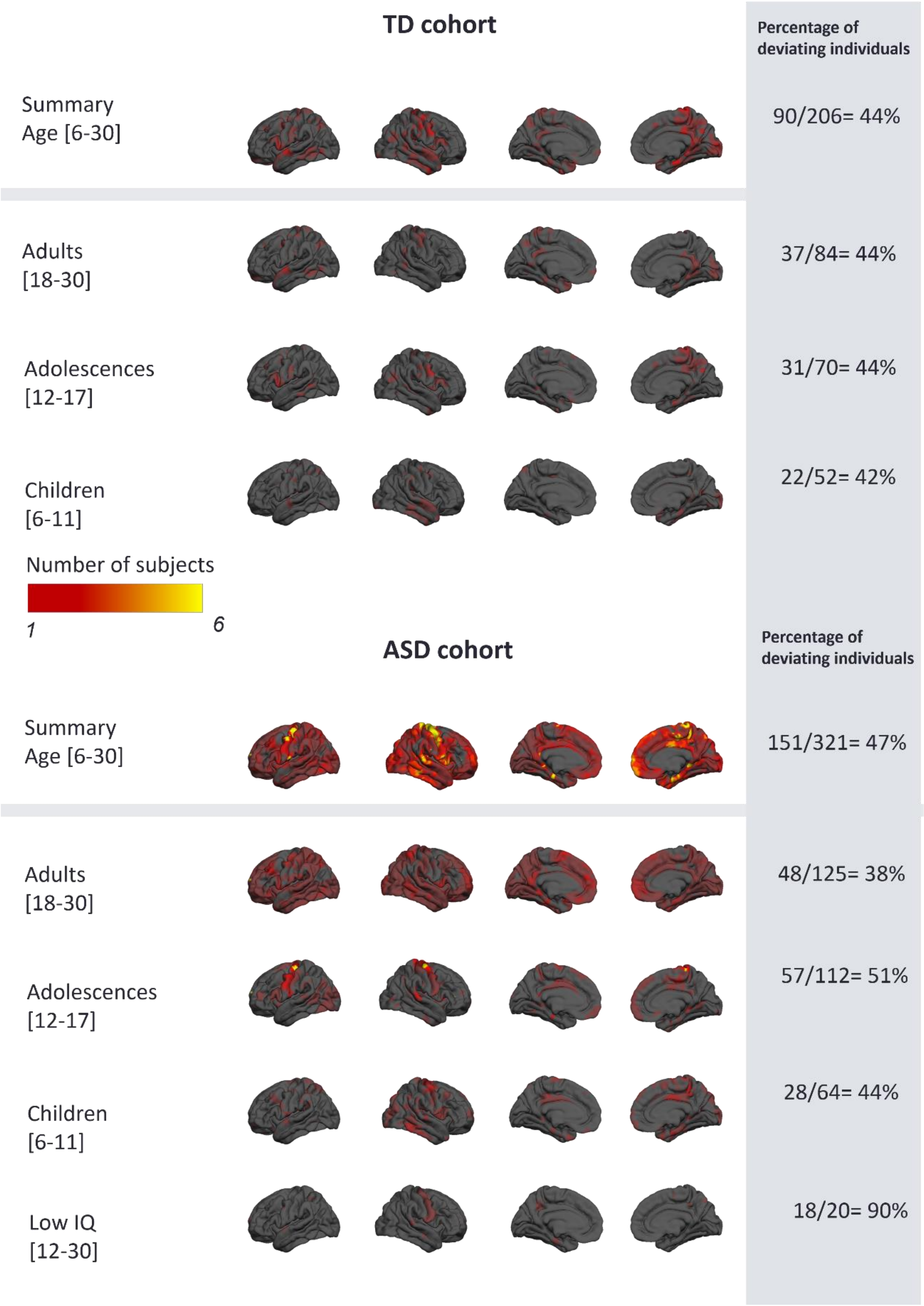

### Individual subject deviations

Supplementary Figure 8 shows the top 15 subjects deviating from the normative pattern for CT.

*Supplementary Figure 8:  NMPs of top fifteen deviating individuals from normative CT model. These subjects who belong to ASD cohort, have highly individualized patterns of deviation with respect to brain regions and different sign of the deviation*

| EV rank | Age | Sex |
|---|---|---|
| 14 | 6.45 | M |
| 10 | 8.38 | M |
| 1 | 9.69 | M |
| 15 | 9.79 | M |
| 6 | 13.60 | M |
| 4 | 15.24 | M |
| 9 | 15.38 | M |
| 12 | 16.25 | M |
| 8 | 18.24 | M |
| 2 | 22.58 | M |
| 5 | 6.24 | F |
| 11 | 7.88 | F |
| 13 | 10.19 | F |
| 7 | 10.68 | F |
| 3 | 18.66 | F |

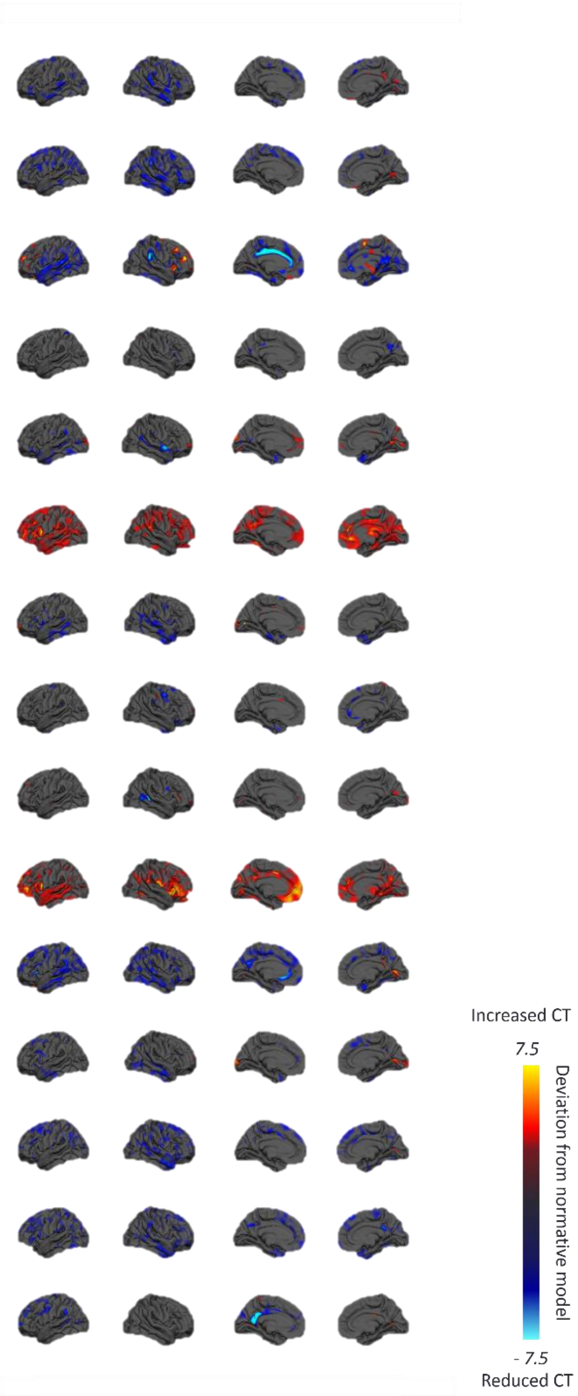

### Individual subject deviations

Supplementary Table 1 shows the clinical characteristics of the sample separately for each site.

*Supplementary Table 1: Clinical characteristics for each site*

| | Cambridge | | *KCL* | | Mannheim | | *Nijmegen* | | *Rome* | | *Utrecht* | |
|---|---|---|---|---|---|---|---|---|---|---|---|---|
| *variable* | *ASD* | *TD* | *ASD* | *TD* | *ASD* | *TD* | *ASD* | *TD* | *ASD* | *TD* | *ASD* | *TD* |
| Age, mean, [SD] | 17.6 [5.8] | 17.0 [6.4] | 16.9 [5.9] | 18.7 [6.6] | 15.4 [3.4] | 15.3 [3.6] | 16.2 [5.6] | 15.0 [4.1] | 24.9 [2.9] | 24.9 [3.5] | 16.8 [5.5] | 16.7 [6.2] |
| IQ , mean [SD] | | | | | | | | | | | | |
| Global IQ | 106 [19] | 114 [11] | 100 [21] | 111 [16] | 102 [13] | 109 [15] | 97 [18] | 101 [14] | 101 [14] | 106 [10] | 105 [13] | 111 [8] |
| Performance IQ | 109 [21] | 116 [12] | 100 [20] | 110 [16] | 104 [15] | 113 [13] | 97 [22] | 102 [18] | 104 [18] | 103 [15] | 107 [17] | 109 [12] |
| Verbal IQ | 103 [17] | 109 [11] | 99 [20] | 111 [18] | 101 [15] | 101 [14] | 97 [19] | 100 [15] | 98 [16] | 109 [8] | 105 [14] | 113 [14] |
| ADI-R [SD] | | | | | | | | | | | | |
| Social | 17.2 [6.6] | - | 17.9 [6.4] | | 15.2 [7.2] | - | 14.4 [6.4] | - | 11.4 [5.7] | - | 16.3 [5.8] | - |
| Communication | 14.5 [2.7] | - | 15.2 [5.4] | - | 10.4 [4.9] | - | 12.7 [5.4] | - | 9.3 [5.4] | - | 11.3 [5.4] | - |
| Repetitive Behavior | 5.0 [2.7] | - | 5.0 [2.4] | - | 5.1 [3.8] | - | 2.9 [2.1] | - | 5.3 [2.3] | - | 3.5 [2.7] | - |
| ADOS [SD] | | | | | | | | | | | | |
| Total | 5.2 [2.4] | - | 5.1 [2.9] | - | - | - | 5.3 [2.7] | - | - | - | 4.8 [2.7] | - |
| Social | 6.3 [1.8] | - | 5.3 [2.8] | - | - | - | 6.1 [2.5] | - | - | - | 5.5 [2.7] | - |
| Repetitive Behavior | 4.5[2.5] | - | 5.7 [2.6] | - | - | - | 3.7[2.7] | - | - | - | 4.3[2.4] | - |
| Schedule | | | | | | | | | | | | |
| A: Adults | 16 | 9 | 50 | 33 | 5 | 5 | 23 | 10 | 17 | 10 | 14 | 10 |
| B: Adolescence | 15 | 6 | 34 | 18 | 20 | 11 | 31 | 27 | 0 | 0 | 12 | 8 |
| C. Children | 7 | 7 | 26 | 9 | 3 | 5 | 20 | 18 | 0 | 0 | 8 | 13 |
| D. IQ < 70 | 2 | - | 12 | - | 0 | - | 6 | - | 0 | - | 0 | - |

#### Scan quality associations with extreme deviations and age

In Supplementary Table 2, we show the correlation of deviations with the Freesurfer Euler number. Note that smaller EN values are indirectly associated with a lower scan quality.

*Supplementary Table 2: correlation between extreme deviations and Euler number*

| | Euler number in ASD cohort | Euler number in TD cohort | Euler number overall |
|---|---|---|---|
| extreme deviations | -0.57 * | -0.62 * | -0.58 * |
| age | 0.39 * | 0.38 * | 0.38 * |
| ADI Social | -0.18 * | - | - |
| ADI Communication | -0.25 * | - | - |
| ADI RRB | -0.25 * | - | - |
| ADOS Total | -0.11 | - | - |
| ADOS Social | -0.02 | - | - |
| ADOS RRB | -0.21 * | - | - |

This shows that EN was correlated with the deviations in both ASD and TD cohorts, all ADI symptom domains, and ADOS repetitive behaviors. Regarding alternative potentially confounding variables, the extreme value deviations were weakly negatively correlated with IQ in the ASD cohort ($\rho$ = -0.16, p<0.05), but not in the TD cohort ($\rho$ = 0.08, n/s). They were also correlated with some ADHD symptom scales (Supplementary Table 3).

*Supplementary Table 3: correlation between extreme deviations and ADHD score*

| | ADHD Inattentive parent in ASD | ADHD Inattentive parent in TD | ADHD Inattentive |
|---|---|---|---|
| extreme deviations | 0.20 * | -0.01 | 0.22 * |
| | ADHD hyperimpulsive parent in ASD | ADHD hyperimpulsive parent in TD | ADHD hyperimpulsive |
| extreme deviations | 0.24 * | 0.16 | 0.26 * |

Taken together these results reinforce the cautions noted above, and preclude a definitive assessment of the degree to which any one particular confounding variable may have influenced our results.

Supplementary Table 4 shows the clinical characteristics from the top deviating subjects from the normative model for CT, all of whom have ASD

*Supplementary Table 4: Clinical characteristics of the top fifteen deviating participants from normative CT model. These subjects who belong to ASD cohort, have highly individualized patterns of deviation with respect to brain regions and different sign of the deviation*

| | | *Schedule* | *Site* | *Sex* | *VIQ* | *PIQ* | *FSIQ* | *Age* | *ADI-social* | *ADI-communication* | *ADI-RRB* | *ADOS-TOTAL* | *ADOS-SA* | *ADOS-RRB* |
|---|---|---|---|---|---|---|---|---|---|---|---|---|---|---|
| 1 | ASD | Children | Mannheim | M | - | 69 | - | 9.69 | 20 | 10 | 8 | - | - | - |
| 2 | ASD | Adults | Rome | M | 70 | 79 | 76 | 22.58 | 18 | 11 | 3 | - | - | - |
| 3 | ASD | Adults | Nijmegen | F | 79 | 66 | 74 | 18.66 | 4 | 6 | 0 | 3 | 4 | 6 |
| 4 | ASD | Adolescence | Utrecht | M | 86 | 106 | 96 | 15.24 | 10 | 9 | 2 | 5 | 6 | 6 |
| 5 | ASD | Children | KCL | F | 90 | 109 | 100 | 6.24 | 17 | 17 | 7 | 10 | 10 | 8 |
| 6 | ASD | Adolescence | Nijmegen | M | 114 | 124 | 118 | 13.6 | 15 | 15 | 2 | - | - | - |
| 7 | ASD | Children | KCL | F | 116 | 133 | 127 | 10.68 | 10 | 9 | 8 | 1 | 2 | 6 |
| 8 | ASD | Adults | KCL | M | 87 | 86 | 84 | 18.42 | - | - | - | 2 | 2 | 6 |
| 9 | ASD | IQ < 70 | KCL | M | 58 | 59 | 56 | 15.83 | 22 | 21 | 9 | 10 | 10 | 10 |
| 10 | ASD | Children | KCL | M | 111 | 106 | 110 | 8.38 | 16 | 18 | 5 | 6 | 3 | 9 |
| 11 | ASD | Children | Utrecht | F | 93 | 98 | 95 | 7.88 | 23 | 17 | 3 | 7 | 5 | 9 |
| 12 | ASD | IQ < 70 | Nijmegen | M | 52 | 50 | 54 | 16.25 | 27 | 21 | 6 | 10 | 10 | 8 |
| 13 | ASD | Children | KCL | F | - | - | - | 10.19 | 0 | 8 | 2 | 6 | 7 | 6 |
| 14 | ASD | Children | KCL | M | 99 | 107 | 104 | 6.45 | 14 | 12 | 5 | 2 | 3 | 1 |
| 15 | ASD | Children | Cambridge | 'M | 106 | 128 | 119 | 9.79 | 16 | 15 | 2 | 4 | 5 | 6 |

Supplementary Table 5 shows the clinical characteristics from the top deviating subjects from the normative model for SA.

*Supplementary Table 5: Clinical characteristics of the top fifteen deviating participants from normative surface area model. While 80% of the individuals belong to ASD cohort with highly heterogeneous profiles, there are several individuals in the list who belong to TD cohort.*

| | | *Schedule* | *Site* | *Sex* | *VIQ* | *PIQ* | *FSIQ* | *Age* | *ADI-social* | *ADI-communication* | *ADI-RRB* | *ADOS-TOTAL* | *ADOS-SA* | *ADOS-RRB* |
|---|---|---|---|---|---|---|---|---|---|---|---|---|---|---|
| 1 | ASD | Adults | Nijmegen | F | 71 | 66 | 70 | 17.49 | 9 | 14 | 2 | 7 | 8 | 1 |
| 2 | ASD | IQ <70 | Cambridge | M | 73 | 66 | 67 | 24.29 | 10 | 9 | 5 | 9 | 8 | 10 |
| 3 | TD | Adults | KCL | M | 142 | 136 | 142 | 23.08 | - | - | - | - | - | - |
| 4 | ASD | Adolescents | Nijmegen | M | 89 | 64 | 78 | 12.07 | 17 | 12 | 1 | 3 | 5 | 1 |
| 5 | TD | Adults | Cambridge | F | 104 | 117 | 111 | 18.26 | - | - | - | - | - | - |
| 6 | ASD | Adolescents | Cambridge | M | 103 | 120 | 113 | 14.53 | 16 | 12 | 1 | 7 | 8 | 5 |
| 7 | ASD | Adolescents | KCL | M | 144 | 134 | 142 | 16.82 | 20 | 12 | 5 | 6 | 6 | 7 |
| 8 | ASD | Children | KCL | F | 98 | 106 | 102 | 9.31 | 25 | 20 | 8 | 8 | 7 | 9 |
| 9 | ASD | Children | KCL | F | 116 | 133 | 127 | 10.68 | 10 | 9 | 8 | 1 | 2 | 6 |
| 10 | TD | Adults | Utrecht | M | 105 | 85 | 96 | 22.06 | - | - | - | - | - | - |
| 11 | TD | Children | Cambridge | M | 108 | 129 | 120 | 8.62 | - | - | - | - | - | - |
| 12 | ASD | Children | Mannheim | M | - | 69 | - | 9.69 | 20 | 10 | 8 | - | - | - |
| 13 | ASD | Adults | KCL | M | 133 | 120 | 130 | 19.44 | 18 | 19 | 8 | - | - | - |
| 14 | ASD | Adolescents | Cambridge | F | 112 | 104 | 109 | 12.11 | 27 | 17 | 8 | 2 | 4 | 1 |
| 15 | ASD | Children | Nijmegen | M | 92 | 94 | 93 | 11.57 | 24 | 20 | 7 | 1 | 3 | 1 |

Finally, Supplementary Tables 6 and 7 shows the associations for the deviations from the normative model and symptom scales.

*Supplementary Table 6: Clinical relevance of the deviations; Significant correlation (Spearman) between the mean of extreme deviation in each cortical parcel and symptoms measured by ADOS and ADI scores (P_value < 0.05). * indicates the regions survived after FDR correction*

| Parcel | Correlation Coefficient, r | Parcel | Correlation Coefficient, r |
|---|---|---|---|
| *ADI_social, Female* | | *ADI_RRB, Male* | |
| superiortemporal | 0.22(L) | superiorfrontal | 0.23*(L), 0.2(R) |
| lateralorbitofrontal | 0.22(R) | superiortemporal | 0.16(L) |
| parsopercularis | 0.22(R) | insula | 0.16(L) |
| precuneus | 0.24(R) | caudalanteriorcingulate | 0.2(R) |
| temporalpole | 0.26(R) | fusiform | 0.15(R) |
| *ADI_social, Male* | | paracentral | 0.15(R) |
| superiortemporal | 0.17(L) | parstriangularis | 0.17(R) |
| lateralorbitofrontal | 0.15(R) | temporalpole | 0.16(R) |
| *ADI_communication, Female* | | *ADOS_II_CSS, Female* | |
| caudalmiddlefrontal | 0.21(L) | parsopercularis | 0.25(L) |
| lateralorbitofrontal | 0.25(L) | *ADOS_II_CSS, Male* | |
| parsopercularis | 0.22(L) | entorhinal | 0.18(R) |
| pericalcarine | 0.23(L) | medialorbitofrontal | 0.19(R) |
| postcentral | 0.22(L) | temporalpole | 0.16(R) |
| precentral | 0.23(L) | *ADOS_II_Social Affect, Female* | |
| superiorparietal | 0.23(L) | entorhinal | -0.23(L) |
| inferiortemporal | 0.23(R) | lateraloccipital | -0.25(L) |
| lateraloccipital | 0.30(R) | lingual | -0.22(L) |
| middletemporal | 0.30(R) | middletemporal | -0.24(L) |
| precuneus | 0.21(R) | parsopercularis | 0.23(L) |
| rostralmiddlefrontal | 0.21(R) | *ADOS_II_Social Affect, Male* | |
| *ADI_communication, Male* | | medialorbitofrontal | 0.18(R) |
| superiortemporal | 0.17(L) | temporalpole | 0.15(R) |
| supramarginal | 0.15(L) | *ADOS_II_RRB, Female* | |
| insula | 0.17(L) | caudalmiddlefrontal | 0.33*(L), 0.31(R) |
| superiorparietal | 0.15(R) | entorhinal | 0.23(L) |
| paracentral | 0.15(R) | fusiform | 0.33*(L) |
| *ADI_RRB, Female* | | inferiorparietal | 0.26(L),0.25(R) |
| caudalmiddlefrontal | 0.31*(L), 0.24(R) | inferiortemporal | 0.25(L) |
| entorhinal | 0.24(L) | lateraloccipital | 0.23(L) |
| inferiortemporal | 0.22(L), 0.28(R) | lingual | 0.38*(L),0.34(R) |
| lateraloccipital | 0.30*(L) | rostralmiddlefrontal | 0.23(L),0.31(R) |
| lateralorbitofrontal | 0.32*(L) | superiorfrontal | 0.33*(L),0.24(R) |

| | | | |
|---|---|---|---|
| medialorbitofrontal | 0.26(L), 0.24(R) | superiorparietal | 0.25(L) |
| parsopercularis | 0.28(L) | supramarginal | 0.33(L) |
| parstriangularis | 0.31*(L) | caudalanteriorcingulate | 0.26(R) |
| precentral | 0.25(L), 0.25(R) | middletemporal | 0.24(R) |
| precuneus | 0.22(L) | paracentral | 0.25(R) |
| superiorparietal | 0.24(L) | pericalcarine | 0.26(R) |
| inferiorparietal | 0.24(R) | precentral | 0.29(R) |
| middletemporal | 0.32*(R) | precuneus | 0.29(R) |
| rostralmiddlefrontal | 0.28(R) | frontalpole | 0.32(R) |
| superiortemporal | 0.23(R) | *ADOS_II_RRB, Male* | |
| supramarginal | 0.36(R) | lateralorbitofrontal | 0.18(L) |
| *ADI_RRB, Male* | | medialorbitofrontal | 0.21(L) |
| entorhinal | 0.19(L) | rostralmiddlefrontal | 0.16(L) |
| inferiortemporal | 0.14(L), 0.17(R) | insula | 0.16(L) |
| lingual | 0.14(L) | entorhinal | 0.15(R) |
| middletemporal | 0.18(L), 0.14(R) | fusiform | 0.17(R) |
| parahippocampal | 0.14(L) | inferiortemporal | 0.21(R) |
| rostralanteriorcingulate | 0.15(L), 0.16(R) | middletemporal | 0.23(R) |
| rostralmiddlefrontal | 0.15(L) | postcentral | 0.18(R) |

*Table 7: Clinical relevance of the deviations across the whole brain; Significant correlation (Spearman) between extreme value across all the regions and symptoms measured by ADOS and ADI scores (P_value < 0.05). * indicates the regions survived after FDR correction*

| | *ADI* | | | *ADOS* | | |
|---|---|---|---|---|---|---|
| | *ADI-social* | *ADI-communication* | *ADI-RRB* | *ADOS-TOTAL* | *ADOS-SA* | *ADOS-RRB* |
| Female | 0.07 | 0.16 | 0.22 | 0.03 | -0.08 | 0.27 |
| Male | 0.07 | 0.04 | 0.11 | 0.05 | -0.00 | 0.16 |
